# Exploring environmental microfungal diversity through serial single cell screening

**DOI:** 10.1101/2024.05.16.594478

**Authors:** Joana Mariz, Ali Nawaz, Yvonne Bösch, Christian Wurzbacher

**Affiliations:** Chair of Urban Water Systems Engineering, Technical University of Munich, Am Coulombwall 3, 85748 Garching, Germany; Department of Digital Health Sciences and Biomedicine, School of Life Sciences, University of Siegen, Am Eichenhang 50, 57076, Siegen, Germany

**Keywords:** Laser-Microdissection, dark fungal taxa, aquatic hyphomycetes, single cell genomics, ribosomal operon, taxonomy, barcoding, biodiversity

## Abstract

Known for its remarkable diversity and ecological importance, the fungal kingdom remains largely unexplored. In fact, the number of unknown and undescribed fungi is predicted to exceed the number of known fungal species by far. Despite efforts to uncover these dark fungal taxa, we still face inherent sampling biases and methodological limitations. Here, we present a framework that combines taxonomic knowledge, molecular biology, and data processing to explore the fungal biodiversity of enigmatic aquatic fungal lineages. Our work is based on serial screening of environmental fungal cells to approach unknown fungal taxa. Microscopic documentation is followed by DNA analysis of laser micro-dissected cells, coupled with a ribosomal operon barcoding step realized by long-read sequencing, followed by an optional whole genome sequencing step. We tested this approach on a range of aquatic fungal cells mostly belonging to the group of aquatic hyphomycetes derived from environmental samples. From this initial screening, we were able to identify thirty-two potentially new fungal taxa in the target dataset. By extending this methodology to other fungal lineages associated with different habitats, we expect to increasingly characterize the molecular barcodes of dark fungal taxa in diverse environmental samples. This work offers a promising solution to the challenges posed by unknown and unculturable fungi and holds the potential to be applied to the diverse lineages of undescribed microeukaryotes.

## Introduction

Fungi, estimated to comprise between 2.2 – 3.8 million species, have successfully colonized nearly every habitat on earth [Blackwell 2011; Taylor et al. 2014; Hawksworth and Lücking 2017]. Nevertheless, with the formal description of currently 156 000 fungal species (https://www.speciesfungorum.org/; 28 May 2023), the fungal inventory is fragmented and incomplete. In other words, despite the sheer amount of evidence supporting the potential impact of fungi in mediating and altering ecosystem dynamics, more than 90% of the fungal kingdom is unknown to science [Cheek et al. 2020; Bahram and Netherway 2022]. Herein, one of the least characterized ecological groups of fungi are those that thrive in aquatic habitats [Jones et al. 2014].

Like in terrestrial environments, fungi can be detected in most surface and subsurface systems (i.e., lakes, streams, rivers, oceans, and aquifers) that have been sampled. It is well established that fungi are key components of aquatic biodiversity [Ittner et al. 2018]. They are known to play a key role in carbon and nutrient cycling, food web dynamics and energy flow in headwater streams [Gladfelter et al. 2019; Grossart et al. 2019; Ruess and Müller-Navarra 2019; Banos et al. 2020]. Despite their ecological key-functions in aquatic ecosystems, the knowledge of aquatic fugal biodiversity is still limited.

In fungal ecology studies, both classical (cultivation, microscopy and characterization of morphological features) and molecular methods (sequencing of extracted DNA and taxonomic markers) have been widely used for the identification and characterization of fungal species in various habitats [Nikolcheva et al. 2005]. Classical culture-based or microscopy-based methods in mycology studies come with an inherent bias of a cultivation bottleneck toward mostly copiotroph or sterile species [Seena et al. 2010] and limited availability of taxonomic keys [Sinha et al. 2012]. Even though spore morphology and conidiogenesis have been widely used to identify aquatic hyphomycetes species in the past [Alexopoulos et al. 1996; Marvanová 1997], this group of fungi is now believed to have a convergent evolution pattern, with distinct phylogenetic lineages independently adapting over time to similar environmental pressures [Belliveau and Bärlocher 2005]. Therefore, conidial morphology is a limited indicator of fungal diversity and phylogenetic relationships [Belliveau and Bärlocher 2005; Campbell et al. 2006]. With the advent of high throughput DNA sequencing and decreasing sequencing costs in the last two decades, sequencing-based solutions have enabled microbial ecologists and mycologists to access species inventories across diverse environments. These sequence-based inventories have uncovered the existence of a large diversity of unidentified fungal species, the so-called dark fungal taxa 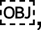 which are still unreported by classical methods [Grossart et al. 2016].

These dark fungal lineages are often found in aquatic habitats [Grossart et al. 2019], resulting in many unclassified fungal barcodes (e.g. Wurzbacher et al. 2016). Aquatic fungi span all fungal lineages, including poorly studied zoosporic fungi e.g., Cryptomycota and Chytridiomycota [Richards et al. 2012; Picard 2017; Tedersoo et al. 2018], and the well-known aquatic hyphomycetes, an ecological group consisting of predominantly ascomycetous fungi [Shearer et al. 2007] belonging to the fungal classes of Leotiomycetes, Sordariomycetes, Dothideomycetes, Orbiliomycetes and Pezizomycetes [Belliveau and Bärlocher 2005; Baschien et al. 2006; Campbell et al. 2006; Campbell et al. 2009; Calabon et al. 2023]. Despite the recognized importance of the aquatic hyphomycetes as the major microbial decomposers of plant litter in streams [Gessner et al. 2007], the taxonomic positioning of many species remains undiscovered. The vast majority of the existing molecular data is constricted to the ITS region, the formal barcode for molecular identification of fungal DNA [Schoch et al. 2012]. Yet, for aquatic hyphomycetes, only for about 42% of described species (136 out of approximately 323 described species) have an ITS barcode sequence listed in GenBank [Franco-Duarte et al. 2022]. Although ITS sequence data has been key to the large advancement of mycological discoveries, there are significant drawbacks when selecting this region as a sole genetic barcode, making it often not adequate to study the broth of fungal lineages. Some of the major adversities are: (i) low taxonomic resolution for species determination (due to rapid species evolution) [Vrålstad 2011; Bates et al. 2013] and (ii) high variability of the region, making it difficult to analyze the sequence data through alignment and phylogeny-based methods across orders [Gazis et al. 2011; Lindner et al. 2011]. The identification of new clades within the dark fungal lineages, such as Cryptomycota, depends on sequence alignment and phylogenetic analysis. Therefore, SSU and LSU regions [Lazarus and James 2015; Burgaud et al. 2016], despite being less variable than the ITS [Porras-Alfaro et al. 2014], have been used for these lineages. One solution that combines the previous efforts on using taxonomic markers is the generation of long-read sequences that cover all of the most commonly used genetic markers for fungi (SSU, ITS-5.8S, and LSU regions). This would allow us to fill many taxonomical and marker-related gaps in our reference sequence databases [Wurzbacher et al. 2019] whilst removing some of the typical constraints of using shorter barcoding regions.

Considering the limitations and challenges of both classical and modern methods employed in mycology so far, in this work, we aim to provide a framework to explore and study the vast universe of so far enigmatic aquatic fungi that can also be transferred to other eukaryotic lineages. Here, we would like to extend the single cell genomics platform recently introduced by Ciobanu and colleagues [Ciobanu et al. 2022] to all fungal (and potentially eukaryotic) cells by linking classical microscopy species identification with modern DNA-based methods on a single cell/spore/conidia level. As a proof of concept, we applied a series of steps from microscopy, laser microdissection of fungal single-cells or mitospores, followed by a whole genome amplification, that served as the basis for taxonomic informative DNA ribosomal tandem repeat barcoding (unifying the most frequently used fungal barcodes: SSU, ITS, LSU), and as a pre-screening for single-cell genomics. The workflow can be handled with relatively short turnaround times in multi-well plate formats allowing for the identification of single cells/spores of various aquatic fungi, simultaneously generating future genetic and morphological reference sequences resources. We tested this framework with fungal cells from pure cultures and environmental samples from different origins. The fungal structures from aquatic samples were **a)** prepared for laser microdissection, **b)** screened and documented under the microscope, **c)** laser-dissected into clean microtubes, **d)** genome amplified using multiple displacement amplification, **e)** screened by PCR ribosomal marker gene amplification and **f)** long-read barcoded by amplicon sequencing, and **g)** whole genome sequenced. In this study, we displayed how this workflow can be used to directly document and barcode unknown fungal cells and extract their genome for further downstream processing. We furthermore tested if the length of the PCR amplicons after the whole genome amplification can be used as a proxy to predict the quality of the obtained genome.

## Results

Here, we present a comprehensive framework that allows barcoding single cells or conidia, by extracting their DNA with a subsequent molecular analysis from aquatic environmental samples. This single cell genomics platform is based on laser dissection microscopy and thus integrates visual inspection, precise dissection, collection, targeted amplification, and sequencing of ribosomal DNA (*rrn*: SSU, ITS-5.8S, and LSU), as well as the option to sequence genomes (see graphic summary in Figure 1). The steps of this workflow include (i) sample collection and pre-processing, (ii) sample preparation for laser microdissection, (iii) microscopic imaging and single-cell dissection, (iv) lysis and whole genome amplification (WGA) of single cells, (v) sequencing of fungal ribosomal barcodes and (vi) amplicon sequence processing and phylogenetic analysis, with the optional additional step of (vii) whole genome sequencing of single cells and sequence processing. The entire workflow (i-vi) can be completed within ten days. A detailed hands-on protocol describing each processing step is provided in the Supplementary Information (SI-Detailed protocol).

**Figure 1:**
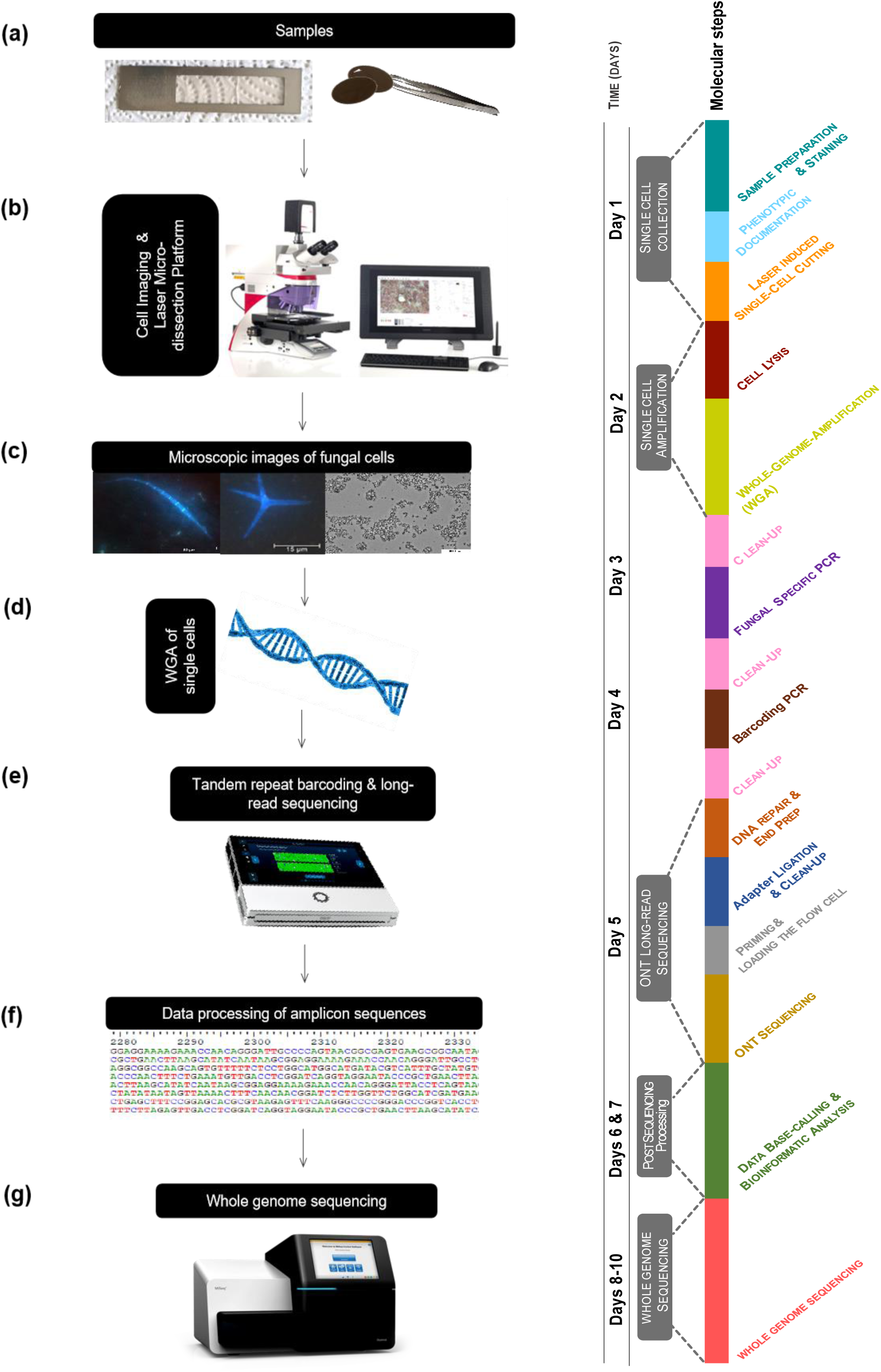
Overview of the methodological framework to identify and resolve the taxonomy and phylogeny of aquatic fungi. **(a)** Fungal cells are concentrated on membrane filters or directly applied to membrane slides **(b)** and placed in the Laser Dissection Microscope. **(c)** The images of the fungal cells are acquired and documented, and the single cells, spores, or conidia are cut and collected in the collection tubes **(d)** for whole genome amplification using multi-displacement amplification (MDA). **(e)** PCR amplification of fungal barcoding regions of amplified single cells (MDA product) and subsequent amplicon sequencing on the MinION (Oxford Nanopore Techonologies) platform. The obtained sequence data is pre-processed to generate fungal short (ITS) and long (FRO) sequences, and the consensus sequences undergo **(f)** further processing by multiple alignments and phylogenetic analysis. Finally, **(g)** whole genome sequencing is performed with using an Illumina platform.

### Barcoding single cells from environmental samples

After initial experimentation with fungal cells from pure cultures, we tested the potential of the workflow for environmental samples and screened several freshwater-derived fungal cells after conidia retrieval (SI-Supplementary data: Table S1). The cells, stained with calcofluor white, were screened for fungal structures and photographed, and their dimensions were measured before targeted laser dissection (Figure 3). Overall, we screened 420 environmental fungal cells/conidia with a dissection efficiency of 65% (Table 1). These successfully dissected cells were lysed based on a combination of enzymatic cell wall digestion and alkaline lysis, followed by a whole genome amplification procedure that served as the basis for barcoding and genomics. We successfully amplified the formal fungal barcode ITS and its extension to the full ribosomal region (*rrn.* operon, FRO) with success rates of 81% and 59%, respectively. Of all dissections attempted from environmental samples, 35% (147 cells/conidia) were successfully barcoded using the extended barcode, only 15% lower than the cells from pure cultures (Table 1), attesting the workflow’s high efficiency for barcoding purposes.

**Table 1:** Overview of the workflow success rates across different work steps: microdissection (cells cut), dissection recovery (cells fallen), ITS amplification (ITS PCR), and FRO amplification (FRO PCR). N: number of tested fungal cells per work step; %: Percentage of successfully treated cells per step.

| Sample type |  | Cells cut | Cells fallen | ITS PCR | FRO PCR |
| --- | --- | --- | --- | --- | --- |
| Pure cultures | Nº | 40 | 30 | - | 22 |
|  | % | - | 75% | - | 73.3% |
| Environmental Samples | Nº | 420 | 274 | 257 | 147 |
|  | % | - | 65.2% | 81.4% | 58.7% |

After sequencing a selection of the barcodes from long (FRO, n = 32) and short (ITS, n = 97) amplicons, we obtained the respective consensus sequences for each type of conidia (SI-Supplementary data: Table S2). Here, we could observe that the majority of ITS sequences (61.5%) resulted in more than one consensus sequence, while only 35% of the FRO sequences showed a secondary consensus sequence. Moreover, putative environmental false positives regarding the dominant consensus sequence for the ITS amplicons were more abundant for the short region (42% vs. 9% for FRO; SI-Supplementary data: Table S3).

### Evaluation of single cell genomes

Despite effectively amplifying both FRO and ITS amplicons in 147 cells, in 110 cells, only the ITS region could be successfully amplified (Table 1). Based on this observation, we assessed if the successful amplification of the entire fungal operon (FRO) was related to DNA integrity determined by gel-electrophoresis and the completeness of genomes obtained from cells for which we performed the whole genome sequencing approach.

We selected a subset of cells representative of samples with and without successful FRO amplification and high DNA integrity for whole genome sequencing and evaluated the subsequent genomes for their completeness. Among the eleven tested whole genome amplified cells, two genomes achieved BUSCO completeness values of over 80%. All other fungal cells exhibited a completeness of about 10.5 ± 3.5 % (mean, SD, n = 9) and an average proportion of 61.4 ± 6.8 % (mean, SD, n = 9) contaminating bacterial sequences. This suggests that bacterial contamination is the main predictor for successful whole genome sequencing of our single cell workflow (SI-Supplementary data: Table S4). To account for bacterial contamination at an early process stage, we implemented an optional quality control measure (viii) in our genome sequencing workflow consisting of a 16S rRNA quantification screening by quantitative PCR (SI-Detailed protocol).

### Identification of dark and known fungal taxa from aquatic environments

For the ITS barcode, we could identify 67 fungi at the species level, out of which 56 cases matched with the morphological identification using taxonomic keys (SI-Supplementary data: Table S2). Molecular identification of the FRO region achieved a species-level identification of seven single cells (which all morphologically and molecularly identified as *Tetracladium marchalianum*). Thirty-two fungal cells from the environmental samples could not be identified at the species level, neither by morphological means nor by ITS or FRO barcodes. These “dark” fungal taxa were subsequently identified by phylogenetic inference using the FRO region, which allowed us to place our single cells into a phylogenetic context using maximum-likelihood trees (Figure 2). Furthermore, the results show taxonomic distribution between 3 phyla (Basidiomycota, Chytridiomycota, and Ascomycota), 9 classes, and 19 families. Most of our sequences belonged to the order Helotiales (49,4% and 46,1% for ITS and FRO barcodes, respectively). The phylogenetic inference resolved the taxonomical characterization of dark fungal taxa to the order level (in the case of the ITS sequences: 46S, 56S, 130S_2, 145S_2). Furthermore, two sequences, 130S_2 and 145S_2 (Chytridiomycetes), obtained from two putative aquatic hyphomycetes conidia, revealed chytrid barcodes for both the ITS and FRO regions. Thus, we suspect that the conidia may have been parasitized with chytrids, representing a first indication that chytrids may use aquatic hyphomycetes as hosts.

**Figure 2:**
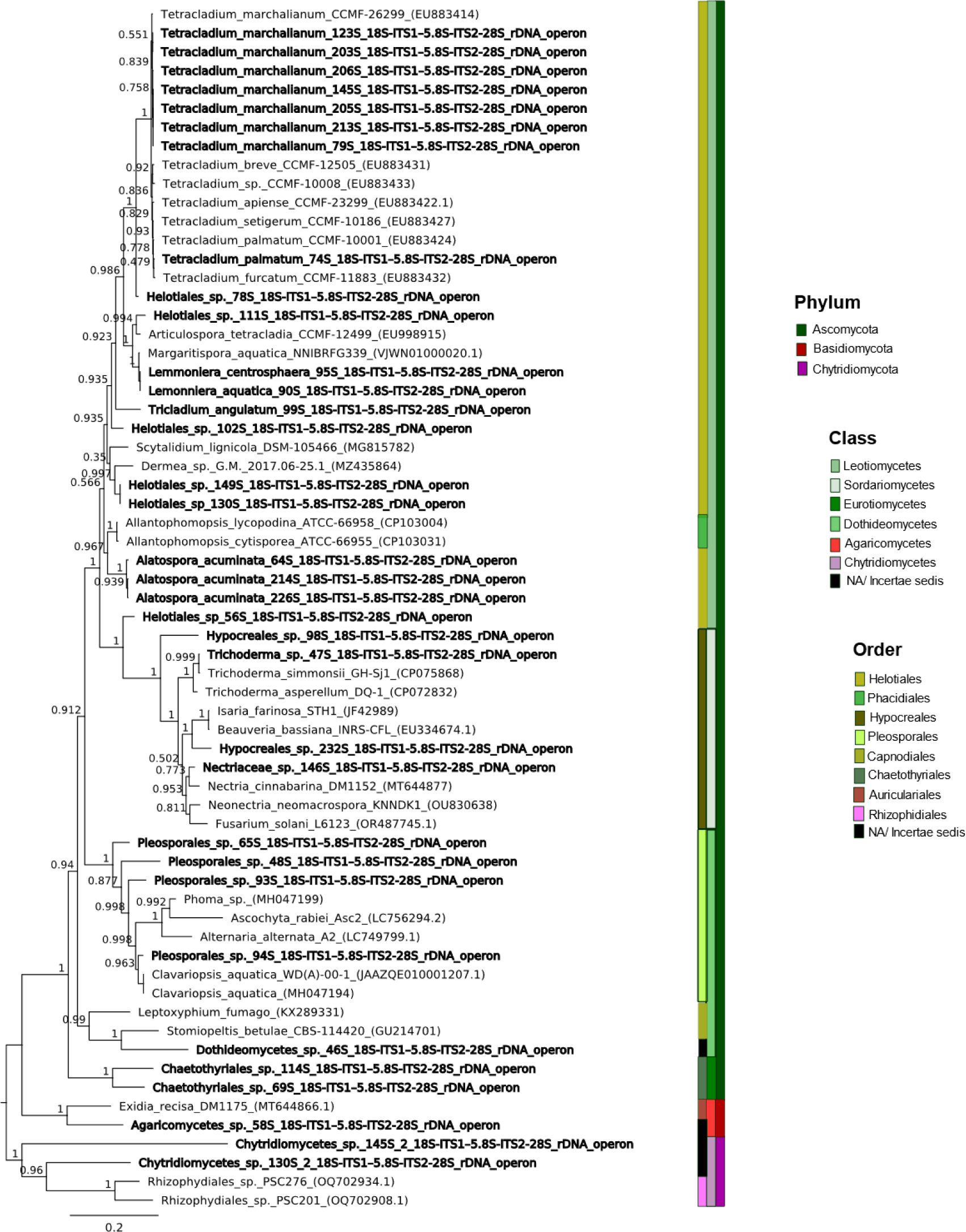
Maximum likelihood phylogenetic tree based on complete fungal ribosomal operon (SSU-ITS1-5.8S-ITS2-LSU) of fungal rDNA. The tree was rooted with the Chytridiomycota branch (sequences PP386605 and PP386607). Chytridiomycota sequences were obtained from the long amplicon sequencing results of samples 130S and 145S. Taxa in bold represent species obtained from environmental samples with the described single cell dissection workflow.

**Figure 3:**
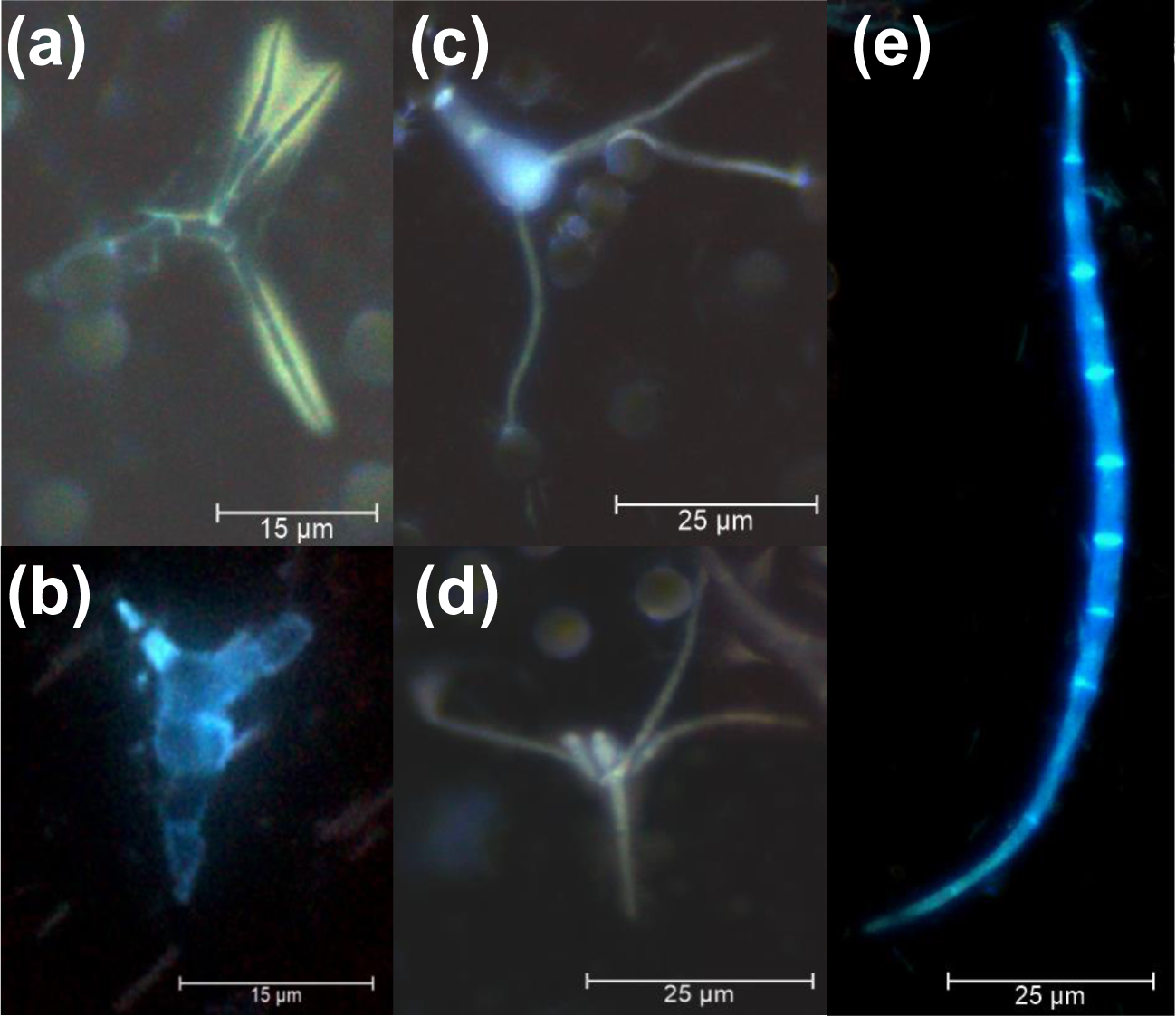
Distinct morphology of conidia obtained from environmental samples at different magnifications. (**a-b**) Environmental conidia (*Alatospora flagellata* and *Chaetothyriales* sp., respectively, observed with an LMD 7 at 600x magnification. (**c-e**) Environmental conidia (*Clavariopsis aquatica, Tetracladium marchalianum* and *Amniculicola longissima,* respectively), 400x).

## Discussion

The coupling of microscopy with molecular biology on the genome scale enables genomics and potential other single cell based molecular analysis of dark fungal taxa and thorough documentation of cells from the environment. By fusing taxonomical knowledge, molecular biology, and bioinformatics, this workflow potentiates advancements in the exploration of fungi and other microeukaryotes. Furthermore, whole genome amplification allows the long-term storage of DNA for each of the dissected cells for later genomic analysis or other molecular biology procedures. The workflow theoretically allows the discovery and dissection of the cells automatically via fluorescence labeling and automatic detection of shapes. However, a manual inspection by a specialist (mycologist, plankton analyst) is still the most efficient way to identify new, non-redundant species and saves resources of the whole genome amplification step.

### Methodological considerations

While laser dissection microscopes are long known to be able to dissect even very small cells such as bacteria [Klitgaard et al. 2005; Gloess et al. 2008], the microdissection process itself requires a high degree of optimization. There are a series of factors that can affect dissection efficiency, namely the room and setup conditions, laser parameters, specimen preparation (like the UV pre-treatment) membrane thickness, sample thickness, and the degree of sample hydration. When setting up this workflow for fungal cells, we first had to establish a clean working environment to avoid contaminations from, e.g., human skin cells. These included enclosing the microscope and the workplace and installing UV treatment steps. Secondly, as fungal cell walls are often rather thick, the recovery of high yields of genomic DNA using conventional extraction methods, such as, for instance, the routine alkaline lysis applied to bacterial single cells is, often times, not feasible [Maaroufi et al. 2004; Rinke et al. 2014]. For this reason, we tested several types of lysis protocols using a a) physical lysis procedure in which we punctured the tip of a cell with the laser, then b) a thermal heat shock for 1 min at 98°, and finally c) the enzymatic treatment with Zymolyase, a mixture of four different enzymes for cell wall degradation [Phalip et al. 2004; Fuxman B. et al. 2016], which eventually worked best (in terms of successfully obtaining the long FRO fragment) in combination with the downstream alkaline lysis from the WGA step. However, aquatic fungi are rarely highly melanized, and therefore, it has not yet been possible to test the lysis efficiency of melanized fungal cells.

The identification of fungal cells and spores was facilitated by the staining of the cells with a fluorescence dye (similar to planktonic samples that are screened for micromanipulation [van den Wyngaert et al. 2022; Nakanishi et al. 2023]) and by using polycarbonate filter membranes, which work well with the equipped laser. The laser shapes followed the cell morphology as closely as possible to minimize contamination. However, we still observed a certain degree of bacterial and fungal contaminants, which may be related to sample concentration on a membrane filter, which has the constraint of enhancing the likelihood of having multiple cells on top or in close proximity to each other, cell fragments, and adsorbed environmental DNA (eDNA); although this is generally considered to be low for polycarbonate filters [Liang and Keeley 2013]). Consequently, a step targeting the removal of bacterial and eDNA contaminants by introducing additional washing steps with low-salt solutions, for example, will be implemented in the future.

Moreover, the status of the cell itself influences the MDA success rate, as it could be a dead cell in which we would expect a higher degree of DNA fragmentation. This may also explain why the success rates for ITS amplification of the environmental samples were higher than for the FRO, as short environmental DNA targets could be co-amplified from fungi derived from the conidia enrichment incubations. Remarkably, the majority of these observed putative contaminants are described as saprobes, commonly found in decaying vegetation (*Purpureocilium lilacinum*, [Luangsa-Ard et al. 2011]), coniferous litter (*Chalara piceae-abietis,* [Koukol 2011]) and pine needles (*Sporidesmium goidanichii*, [Tokumasu 2001]). For the present case of *Tumularia aquatica,* a known aquatic hyphomycete, there were no accounts of its presence in our samples during the image acquisition and documentation stage of the workflow. However, we obtained multiple ITS consensus sequences of this species in our samples. These were likely introduced during the 24-hour aeration of the leaves when fungal fragments and DNA from the leaf litter were distributed in the water. Interestingly, the FRO of these putative ITS contaminants did not exhibit matches to these species, pointing to more fragmented environmental DNA. This emphasizes the use of the ribosomal operon for single cell identification, as it allows us to use this as an effective pre-screening for intact DNA from the target cell of an environmental sample.

In this work, our data could not be used to test the hypothesis that the successful amplification of long barcodes indicates whole genome integrity and quality, as the bacterial load was the dominant predictor. However, to avoid contaminants from fungal environmental DNA, cells with an FRO barcode are a first and important quality criterion that will simultaneously allow us to place the unknown fungal cell into a phylogenetic context.

Microbial ecologists have recently been able to adapt and combine a variety of tools previously developed in far-ranging fields [Grujcic et al. 2022] for fungal identification, such as matrix-assisted laser desorption/ionization time-of-flight mass spectrometry (MALDI-TOF MS) [Cornut et al. 2019]; single-spore isolation, followed by marker gene (rDNA) barcoding sequencing [Bärlocher et al. 2010; Karpov et al. 2020; Kagami et al. 2021; Strassert et al. 2021; van den Wyngaert et al. 2022; Nakanishi et al. 2023] and even single-spore isolation, genome amplification and whole-genome sequencing [Ciobanu et al. 2022]. While the single cell analysis using flow cytometry is an established method, it is limited by the size of the cells and only works in an automated manner, which usually excludes manual inspections by cells. Imaging flow cytometry has integrated microscopy-based optics, and thus, some steps of our workflow may be adapted to such an instrument for high throughput. Micromanipulation and single cell picking e.g., [van den Wyngaert et al. 2022; Nakanishi et al. 2023] allows an analogous workflow as the laser microdissection, however, is also accompanied by a high time investment. A laser microdissection microscope, as used in this work, has the advantage of using different magnifications adjusted to the cell’s morphology and size that enables informed decisions on the relevance and integrity of a cell of interest. This also reduces the throughput compared to flow cytometry but increases the quality and accuracy of the cells of interest. Furthermore, the image analysis and documentation can be done in a separate step, which is important for obtaining reference image materials that can be deposited along the barcodes (similar as in SI-Supplementary data: Table S2).

### Accurate classification of aquatic hyphomycetes conidia

Accurately identifying freshwater fungi by morphology, specifically conidia, is a challenging task that requires extensive taxonomical expertise and is associated with a high degree of uncertainty [Bärlocher 1992]. There are several cases where only subtle morphological differences can be observed between species, therefore requiring knowledgeable examinations of the conidiogenesis processes, conidiophoric structures, as well as multiple conidia in order to establish unambiguous species identifications (e.g. [Roldán et al. 1989]). For example, sigmoid (worm-like) spores are observed for several genera of aquatic ascomycetes [Webster and Davey 1984], thereby impeding unambiguous taxonomic identification [Voglmayr 1996; Garnett et al. 2000]. In our study, a proportion of more than one-third of the cells showed sigmoid morphologies. Hence, using a molecular barcode for the identification of less represented taxa among sigmoid cells, as performed in our workflow, has the potential to significantly improve our understanding of aquatic fungal diversity [Bärlocher 2010; Nilsson et al. 2019], taking into account that aquatic hyphomycetes and other aquatic fungi are taxonomically highly divergent [Franco-Duarte et al. 2022][Grossart et al. 2019]. As the ITS region as single marker may not be robust enough to cover this diversity [Kauserud 2023], an extension to the neighboring SSU and LSU regions holds to benefit of a more robust phylogenetic placement, and its reusability in future environmental amplicon studies, which usually depend on either one of these three ribosomal regions.

Similar to previous studies, the ITS region worked as expected for the aquatic hyphomycetes in our samples. As previously reported, the genera *Lemonniera* and *Tetracladium* is represented by a well-defined clade (SI-Supplementary data: Figure S1), however no clear separation among the species of both genera is apparent [Baschien et al. 2013; Franco-Duarte et al. 2022]. Although having less data for the FRO, this pattern seems to persists when considering the ITS, SSU and LSU region, seperately (SI-Supplementary data: Figure S2-S4), or all three regions together (Figure 2). On the other hand, there are several instances where phylogenetic inference based on the complete ribosomal fragment (Figure 2) enables a higher support for the phylogenetic placement of dark fungal taxa. An example would be the placement of the two single cells *Chaetothyriales* sp. 114S and 69S, which placement is not well supported by any of the three regions, and which only have a full bootstrap support in the FRO tree (Figure 2), where they form a completely new clade, which may require a renaming of these organisms in the future. Also, the placement of the unknown *Helotiales* sp. is not consistent between the three regions and only the FRO region enabled a fully supported placement of these single cells. These examples highlight the necessity of long barcodes for accurate species descriptions, in particular for dark fungal taxa.

### The potential of the workflow to identify known and dark taxa

The development and further use of this workflow has the potential to bridge the gaps in the fungal reference sequence databases not only by morphological characterization of previously DNA-documented unknown species but also by discovery, characterization, and documentation of new dark fungal species. In addition, small adjustments to the current workflow would allow the visualization, isolation, and collection of sequencing data for other dark fungal lineages inhabiting distinct environments (e.g., Chytridiomycota and Cryptomycota) as well as other eukaryotic species (e.g., diatoms and large bacteria in biofilms or sediments). Moreover, the laser microdissection could be coupled with single cell cultivation attempts to increase the chances of obtaining new isolates. Analogous to existing single cell platforms for free-living prokaryotes [Rinke et al. 2014] or e.g. also free-swimming fungal zoospores [Ahrendt et al. 2018; Ciobanu et al. 2022], our workflow may help to increase the known fraction of fungal diversity by leveraging single cell and single conidia morphological and genomic characterization. While this is particularly relevant in aquatic systems, this may also be of high importance for other ecosystems that harbor a high degree of dark fungal taxa.

## Material and Methods

### Sample collection and pre-processing

Initial tests were performed with resuspended single cells derived from fungal pure cultures (belonging to the genera: *Candida, Tetracladium, Articulospora, Clavariopsis, Trichoderma, Helicoon*) grown on malt extract agar. The workflow was then further developed towards the analysis of environmental samples from natural aquatic environments. Here, we focused on cells derived from leaf litter samples after aeration (SI-Supplementary data: Table S1). To obtain mitospores (conidia) of leave degrading aquatic hyphomycetes, submerged decomposing leaves from freshwater streams were collected in sterile plastic bags in south Germany (Bavaria) in winter 2021/2022. Fungal sporulation was induced by incubating the leaves at room temperature for 48 h - 72 h with a continuous air supply in deionized water [Baerlocher 2005].

### Sample preparation and cell staining for laser microdissection

Cells were either distributed on PPS-Membrane slides (4.0 µm, Molecular Machines & Industries, Germany), which were pre-treated with UV-irradiation using a UVP Ultraviolet Crosslinker CL-100 for 30 minutes at a dosage of 11.7 J/cm^2^ in an upward position with a 10 cm distance to the UV source or filtered on a filter membrane. Fungal conidia obtained from the leaf litter sporulation assay were trapped on three different membrane filter types: a 5 µm polycarbonate filter (Whatman®, Cytiva, Massachusetts, USA, Durapore®), a 10 µm polyvinylidene difluoride (PVDF) filter (Durapore®, Merck Millipore, Cork, Ireland), and a 12 µm cellulose nitrate filter (Whatman®, Cytiva, Massachusetts, USA). After testing several filter types, polycarbonate membrane filters were most suitable for an efficient laser-based microdissection. Additionally, cells were stained directly on the filter or in suspension by calcofluor white that binds to beta glycosidic cell wall components, such as glucan, chitin, and cellulose [Rasconi et al. 2009] (1 g/L). Eventually, PPS-Membrane slides, or filters, which were fixed to UV-radiated LMD metal frames (Molecular Machines & Industries, Germany), were manually processed by the laser microdissection microscope (LMD7, Leica Microsystems, Wetzlar, Germany).

### Microscopic imaging and single-cell dissection

For contact-free laser microdissection of single cells, the laser, equipped in an LMD7 microscope, was first calibrated using LMD software (Leica Microsystems, Wetzlar Germany, v 8.2.7603), and later the cutting was performed. The cells were cut at a slow speed and optimum power to minimize the cell and DNA damage with the laser beam. For every cutting session, four 0.2 ml tubes (Multiply®-Pro, SARSTEDT, Nümbrecht, Deutschland) or eight collection caps (Optical Flat 8-Cap strips for 0.2 ml tube strips/ plates, BIO RAD, California, USA) with PCR plates (Twin.tec™ 96-Well-PCR-plates, Eppendorf SE, Hamburg, Germany) were used. First, the slides were subjected to an overview scan to find the fungal cells of interest using either the bright field or fluorescence feature of the microscope and marked for laser microdissection at either 200x, 400x, or 630x magnification. After cell dissection, the collection caps were inspected to count the successfully dissected target cells. Cells were imaged at all stages (i.e., before and after laser dissection) in order to allow cell size measurements and documentation.

### Lysis and whole genome amplification (WGA) of single cells

Six µl of Advanced storage buffer (REPLI-g Advanced DNA Single Cell Kit – Qiagen) were transferred to each cell/conidia. Fungal cell walls were digested by adding 5 units of Zymolyase (Zymo Research, CA, USA) and further incubation for 60 min at 37 °C. The Zymolyase was deactivated by an incubation at 65 °C for 10 min. This was followed by the alkaline lysis from the REPLI-g kit using the Advanced Buffer DLB (reconstituted in nuclease free water), 1 M dithiothreitol (DTT), and HCl (Stop solution) following the manufacturer’s instructions, followed by a genome amplification via multiple displacement amplification (MDA), with a prolonged incubation period of 4 h at 30 °C [Stepanauskas et al. 2017].

### Barcoding of single cells

The fungal ITS (ITS5/ITS4) and ribosomal operon sequence (NS1short/RCA95m) was targeted using the primers described in [Gardes and Bruns 2008] as described in the detailed protocol (Supplementary Information: SI-Detailed Protocol). Briefly, the amplification success of the ITS1-5.8S-ITS2 region (0.3-0.7 kb) and 18S-ITS1-5.8S-ITS2-28S rDNA operon (∼ 4-6 kb) were documented by gel electrophoresis, amplicons were purified, PCR-barcoded, pooled, and sequenced with Oxford Nanopore using the Q20+ Early Access Kit (SQK-Q20EA) or the SQK-LSK112 library preparation kits in combination with a R10.4 flow cells (FLO-MIN112; Oxford Nanopore Technologies, United Kingdom). Overall, we aimed for 1,000 – 2,000 reads per cell.

### Generation of consensus sequence

The raw fast5 files were basecalled and converted to fastq files employing the high accuracy basecalling mode using guppy (v6.1.7). To increase the quality, the duplex read pairs were merged using ONT’s duplex tools (https://pypi.org/project/duplex-tools/; v0.2.9). The resulting high-quality fastq files (97-100% accuracy) were processed using the bioinformatic workflow described by Wurzbacher et al., 2019 [Wurzbacher et al. 2019]. In brief, sequences were filtered by length, demultiplexed into individual samples as FASTA files based on their unique barcodes, aligned using MAFFT (v7.397) [Katoh and Standley 2013] with the auto-alignment option and clustered using mothur v1.39 [Schloss et al. 2009] with the Opticlust algorithm. The final consensus sequences for each OTU cluster were generated utilizing Consension (v1.0, https://microbiology.se/software/consension/). The consensus sequences and its OTU variants were inspected for putative PCR artifacts. Secondary OTUs with lower read numbers were discarded. In some cases, the most dominant OTU was not related to the single cell (as identified by the conidia morphology) and were subsequently assorted (SI-Supplementary data: Table S2).

### Phylogenetic analysis

Multiple reference sequences were retrieved by blasting the consensus sequences from the single cells with megaBLAST against the NCBI nucleotide database (BLAST+ v.2.15.0 [Camacho et al. 2009]). The closest neighbors were identified based on coverage and similarity as well as sequence length (only sequences with the expected size or bigger were selected). All sequences were trimmed and primers removed using Geneious Prime 2023.2.1 (https://www.geneious.com) prior multiple sequence alignment running MAFFT v7.490 [Katoh and Standley 2013] with the multiple sequence alignment algorithm FFT-NA-1 (gap opening penalty = 1.53 and offset value = 0.123) implemented in Geneious Prime. The resulting nucleotide alignment was manually checked and curated by removing trailing ends or putative introns that were only present in very few sequences. Additionally, within the FRO alignment, primers (NS8: TCC GCA GGT TCA CCT ACG GA, ITS1: TCC GTA GGT GAA CCT GCG G and ITS4: TCC TCC GCT TAT TGA TAT GC [White 1990], and LR0R: ACC CGC TGA ACT TAA GC and LR7: TAC TAC CAC CAA GAT CT [Vilgalys and Hester 1990]) were used to identify the SSU, ITS1-5.8S-ITS2 and LSU regions of the FRO alignment. These regions were individually extracted to create SSU, ITS1-5.8S-ITS2 and LSU alignments. The curated multiple sequence alignments were used to infer the phylogenetic inferences using maximum-likelihood FastTree (version 2.1.11), using the Generalized Time-Reversible (GTR) transition model to generate approximately maximum-likelihood trees. We used a combination of Geneious Prime and Inkscape (version 1.2.2) to rearrange and annotate the final trees.

### Whole genome sequencing of single cells and sequence processing

The genomic DNA of eleven selected cells was subjected to Illumina short-read sequencing. The WGA amplified DNA was purified with 0.8 v/v AMPure XP beads as described above and approximately 50 ng served as input for a Nextera XT library preparation for Illumina NextSeq or MiniSeq sequencing following the manufacturer recommendations. The resulting fastq files were demultiplexed using the Bcl2Fastq (v2.19.0) software (Illumina) and adapter and low-quality sequences (<Q25) as well as reads shorter than 36 bases were trimmed and removed using Trimmomatic (v0.39) [Bolger et al. 2014] [Bolger et al. 2014]. Quality and complete adapter removal was verified using fastQC [Andrews 2010] before sequence assembly by Spades (v3.15.4) [Bankevich et al. 2012]. The completeness of the genomes was evaluated using the eukaryotic BUSCO reference (eukaryota_odb10) [Manni et al. 2021]. The bacterial contamination was determined using Tiara (version 1.0.3) on contigs greater than 3 kb.

## Supporting information

Detailed Protocol

## Acknowledgments

We would like to thank Heidrun Mayrhofer for technical assistance in the laboratory. The Laboratory for Functional Genome Analysis (LAFUGA) of the Ludwig Maximilian-University Munich is thanked for performing the high-throughput sequencing, and the Leibniz-Rechenzentrum (LRZ) for providing the computational resources. This study was funded by the German Research Foundation (DFG, WU 890/2-1).

## Data availability statement

Consensus sequencing data was deposited under the GenBank accession numbers PP386583-PP386616 for FRO, and PP384220-PP384318 for ITS sequences. WGA short read sequences were deposited under the NCBI BioProject accession: PRJNA1092266. The final alignment for the underlying phylogenetic tree was deposited Zenodo (doi:10.5281/zenodo.10961085).

## Author contributions

JM: Methodology, Validation, Resources, Formal analysis, Investigation, Data Curation, Writing – Original Draft, Writing – Review & Editing, Visualization.

AN: Methodology, Validation, Data Curation, Investigation, Writing – Original Draft, Writing – Review & Editing, Visualization, Supervision, Project administration.

YB: Methodology, Supervision, Formal analysis, Writing – Review & Editing.

CW: Conceptualization, Methodology, Validation, Resources, Writing – Original Draft, Writing – Review & Editing, Supervision, Project administration, Funding acquisition.

