## Supplementary Data for "Exploring environmental microfungal diversity through serial single cell screening"

**Table S1:** Collected environmental leaf litter samples with sampling location and number of dissected cells per sample tested for barcoding.

| Sampling Location | Nr. of cells tested |
|---|---|
| Isar, Garching, Germany | 157 |
| Obernachkanal, Germany | 51 |
| Harrer Graben, Germany | 27 |
| Schleifmühlerlane, Germany | 21 |
| Kranzbach, Germany | 37 |
| Lindenbach, Germany | 9 |
| Walchensee, Germany | 4 |
| Kleine Ammmerquellen, Germany | 13 |
| Kankerbach, Germany | 8 |
| Seinsbach, Germany | 5 |
| Maisingerbach, Germany | 26 |
| Ramsach, Germany | 43 |

**Table S2:** Identification and characterization of diverse fungal species through our polyphasic approach, combining phenotypic microscopic images and sequencing of the complete ITS (ITS1-5.8S-ITS2) and/or FRO (18S-ITS1-5.8S-ITS2-28S) regions of the isolated cells.

| Identifier | Photo | Sequence | | Microscopic identification |
|---|---|---|---|---|
| 9 | 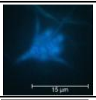 | ITS | PP384220 | *Tetracladium breve* |
| 10 | 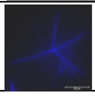 | ITS | PP384221 | *Lemonniera centrosphaera* |
| 32 | 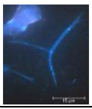 | ITS | PP384222 | *Alatospora acuminata* |
| 37 | 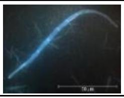 | ITS | PP384223 | *Amniculicola* sp. |
| 38 | 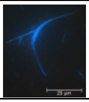 | ITS | PP384224 | *Lunulospora curvula* |
| 40 | 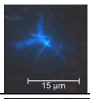 | ITS | PP384225 | *Lemonniera terrestris* |
| 42 | 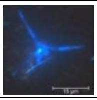 | ITS | PP384226 | *Tetracladium marchalianum* |
| 46 | 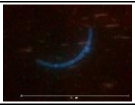 | ITS | PP384227 | *Dothideomycetes* sp. |
| | | FRO | PP386583 | |

| | | | | |
|---|---|---|---|---|
| 47 | 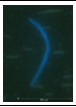 | ITS | PP384228 | *Trichoderma* sp. |
| | | FRO | PP386584 | |
| 48 | 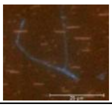 | ITS | PP384229 | *Pleosporales* sp. |
| | | FRO | PP386585 | |
| 50 | 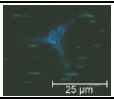 | ITS | PP384230 | *Tetracladium marchalianum* |
| 56 | 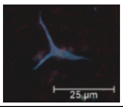 | ITS | PP384232 | *Helotiales* sp. |
| | | FRO | PP386586 | |
| 58 | 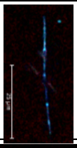 | ITS | PP384233 | *Agaricomycetes* sp. |
| | | FRO | PP386587 | |
| 61 | 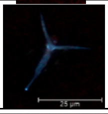 | ITS | PP384234 | *Lemonniera centrosphaera* |
| 64 | 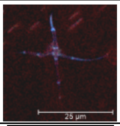 | ITS | PP384235 | *Alatospora acuminata* |
| | | FRO | PP386588 | |
| 65 | 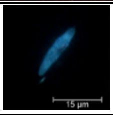 | ITS | PP384236 | *Pleosporales* sp. |
| | | FRO | PP386589 | |
| 69 | 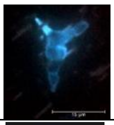 | ITS | PP384237 | *Chaetothyriales* sp. |
| | | FRO | PP386590 | |
| 73 | 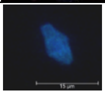 | ITS | PP384238 | *Exophiala cancerae* |
| 74 | 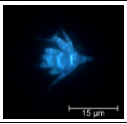 | ITS | PP384239 | *Tetracladium palmatum* |
| | | FRO | PP386591 | |
| 78 | 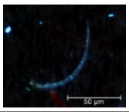 | ITS | PP384240 | *Helotiales* sp. |
| | | FRO | PP386592 | |
| 79 | 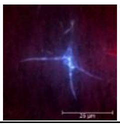 | ITS | PP384241 | *Tetracladium marchalianum* |
| | | FRO | PP386593 | |
| 85 | 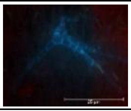 | ITS | PP384242 | *Tricladium angulatum* |
| 90 | 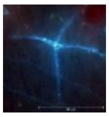 | ITS | PP384243 | *Lemonniera aquatica* |
| | | FRO | PP386594 | |
| 93 | 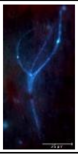 | ITS | PP384244 | *Pleosporales* sp. |
| | | FRO | PP386595 | |

| | | | | |
|---|---|---|---|---|
| 94 | 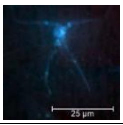 | ITS | PP384245 | *Pleosporales* sp. |
| | | FRO | PP386596 | |
| 95 | 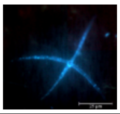 | ITS | PP384246 | *Lemmoniera centrosphaera* |
| | | FRO | PP386597 | |
| 98 | 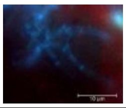 | ITS | PP384247 | *Hypocreales* sp. |
| | | FRO | PP386598 | |
| 99 | 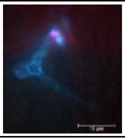 | ITS | PP384248 | *Tricladium angulatum* |
| | | FRO | PP386599 | |
| 102 | 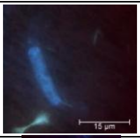 | ITS | PP384249 | *Helotiales* sp. |
| | | FRO | PP386600 | |
| 104 | 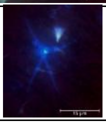 | ITS | PP384250 | *Tetracladium marchalianum* |
| 111 | 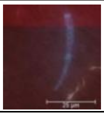 | ITS | PP384251 | *Helotiales* sp. |
| | | FRO | PP386601 | |
| 112 | 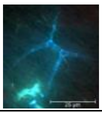 | ITS | PP384252 | *Tricladium angulatum* |
| 114 | 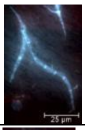 | ITS | PP384253 | *Chaetothyriales* sp. |
| | | FRO | PP386602 | |
| 115 | 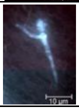 | ITS | PP384254 | *Tetracladium marchalinaum* |
| 123 | 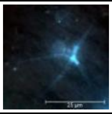 | ITS | PP384255 | *Tetracladium marchalianum* |
| | | FRO | PP386603 | |
| 126 | 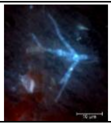 | ITS | PP384256 | *Helotiales* sp. |
| 130 | 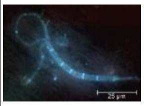 | ITS | PP384257 | *Helotiales* sp. |
| | | FRO | PP386604 | |
| 130_2 | | ITS | PP384258 | *Chytridiomycetes sp.* |
| | | FRO | PP386605 | |
| 133 | 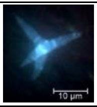 | ITS | PP384259 | *Lemonniera terrestris* |
| 136 | 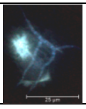 | ITS | PP384260 | *Helotiales* sp. |

| | | | | |
|---|---|---|---|---|
| 142 | 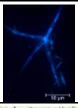 | ITS | PP384261 | *Alatospora acuminata* |
| 143 | 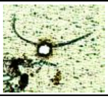 | ITS | PP384262 | *Amniculicola sp.* |
| 145 | 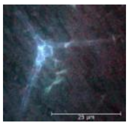 | ITS | PP384263 | *Tetracladium marchalianum* |
| | | FRO | PP386606 | |
| 145_2 | | ITS | PP384264 | *Chytridiomycetes sp.* |
| | | FRO | PP386607 | |
| 146 | 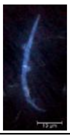 | ITS | PP384265 | *Nectriaceae* sp. |
| | | FRO | PP386608 | |
| 147 | 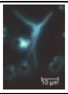 | ITS | PP384266 | *Tetracladium marchalianum* |
| 149 | 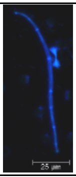 | ITS | PP384267 | *Helotiales* sp. |
| | | FRO | PP386609 | |
| 152 | 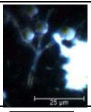 | ITS | PP384268 | *Alatospora acuminata* |
| 154 | 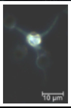 | ITS | PP384269 | *Tetracladium marchalianum* |
| 165 | 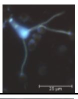 | ITS | PP384270 | *Clavariopsis aquatica* |
| 172 | 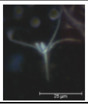 | ITS | PP384271 | *Tetracladium marchalianum* |
| 173 | 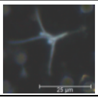 | ITS | PP384272 | *Tetracladium marchalianum* |
| 195 | 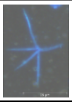 | ITS | PP384273 | *Alatospora acuminata* |
| 202 | 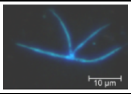 | ITS | PP384274 | *Alatospora acuminata* |
| 203 | 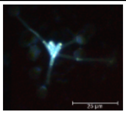 | ITS | PP384275 | *Tetracladium marchalianum* |
| | | FRO | PP386610 | |
| 204 | 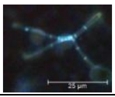 | ITS | PP384276 | *Alatospora acuminata* |
| 205 | 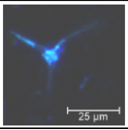 | ITS | PP384277 | *Tetracladium marchalianum* |
| | | FRO | PP386611 | |

| | | | | |
|---|---|---|---|---|
| 206 | 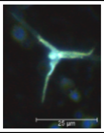 | ITS | PP384278 | *Tetracladium marchalianum* |
| | | FRO | PP386612 | |
| 207 | 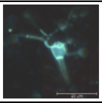 | ITS | PP384279 | *Flagellospora* sp. |
| 209 | 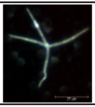 | ITS | PP384280 | *Lemonniera centrosphaera* |
| 210 | 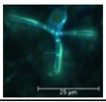 | ITS | PP384281 | *Alatospora acuminata* |
| 211 | 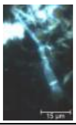 | ITS | PP384282 | *Neonectria lugdunensis* |
| 212 | 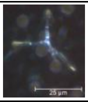 | ITS | PP384283 | *Alatospora acuminata* |
| 213 | 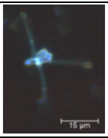 | ITS | PP384284 | *Tetracladium marchalianum* |
| | | FRO | PP386613 | |
| 214 | 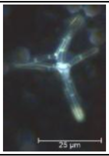 | ITS | PP384285 | *Alatospora acuminata* |
| | | FRO | PP386614 | |
| 215 | 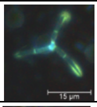 | ITS | PP384286 | *Alatospora acuminata* |
| 216 | 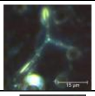 | ITS | PP384287 | *Alatospora acuminata* |
| 218 | 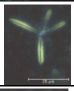 | ITS | PP384288 | *Alatospora acuminata* |
| 219 | 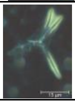 | ITS | PP384289 | *Alatospora flagellata* |
| 220 | 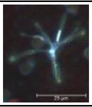 | ITS | PP384290 | *Alatospora acuminata* |
| 221 | 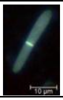 | ITS | PP384291 | *Neonectria lugdunensis* |
| 225 | 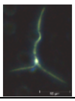 | ITS | PP384292 | *Alatospora acuminata* |
| 226 | 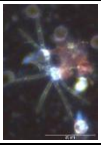 | ITS | PP384293 | *Alatospora acuminata* |
| | | FRO | PP386615 | |

| | | | | |
|---|---|---|---|---|
| 228 | 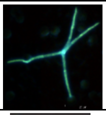 | ITS | PP384294 | *Lemonniera centrosphaera* |
| 231 | 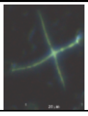 | ITS | PP384295 | *Lemonniera aquatica* |
| 232 | 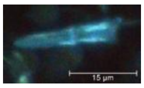 | ITS<br>FRO | PP384296<br>PP386616 | *Hypocreales* sp. |
| 236 | 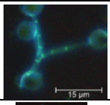 | ITS | PP384297 | *Alatospora acuminata* |
| 247 | 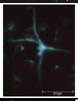 | ITS | PP384298 | *Tetracladium marchalianum* |
| 251 | 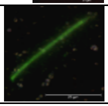 | ITS | PP384299 | *Flagellospora* sp. |
| 253 | 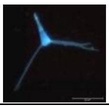 | ITS | PP384300 | *Clavariopsis aquatica* |
| 254 | 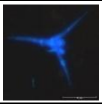 | ITS | PP384301 | *Tetracladium marchalianum* |
| 255 | 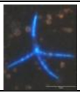 | ITS | PP384302 | *Alatospora acuminata* |
| 257 | 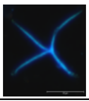 | ITS | PP384303 | *Alatospora acuminata* |
| 258 | 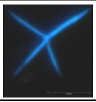 | ITS | PP384304 | *Alatospora acuminata* |
| 259 | 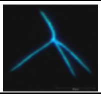 | ITS | PP384305 | *Alatospora flagellata* |
| 261 | 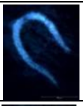 | ITS | PP384306 | *Neoascochyta* sp. |
| 262 | 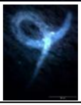 | ITS | PP384307 | *Neoascochyta* sp. |
| 263 | 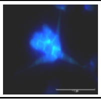 | ITS | PP384308 | *Tetracladium breve* |
| 264 | 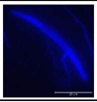 | ITS | PP384309 | *Melanommataceae* sp. |
| 265 | 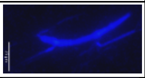 | ITS | PP384310 | *Melanommataceae* sp. |
| 266 | 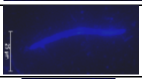 | ITS | PP384311 | *Melanommataceae* sp. |
| 267 | 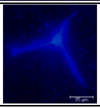 | ITS | PP384312 | *Melanommataceae* sp. |

| | | | | |
|---|---|---|---|---|
| 268 | 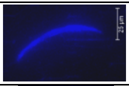 | ITS | PP384313 | *Melanommataceae* sp. |
| 270 | 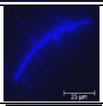 | ITS | PP384314 | *Melanommataceae* sp. |
| 271 | 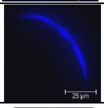 | ITS | PP384315 | *Melanommataceae* sp. |
| 272 | 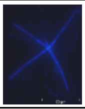 | ITS | PP384316 | *Lemonniera aquatica* |
| 273 | 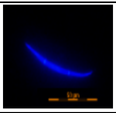 | ITS | PP384317 | *Rhodotorula* sp. |
| 274 | 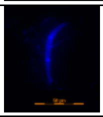 | ITS | PP384318 | *Rhodotorula* sp. |

**Table S3:** Overview of the most common environmental contaminants from leave litter incubations and their prevalence in the results from data processing, identified by the top blast hit.

| **Environmental contaminant (closest BLAST hit)** | **ITS** | **FRO** |
|---|---|---|
| | **OTU1** | **OTU1** |
| *Purpureocilium lilacinum* | 6.6% | ----------- |
| *Chalara piceae-abietis* | 5.5% | ----------- |
| *Sporidesmium goidanichii* | 1.1% | ----------- |
| *Tumularia aquatica* | 13.2% | ----------- |
| Others | 15.4% | 13.5% |
| Total contamination | 41.8% | 13.5% |

**Table S4**: Assessment of assembled genomes of aquatic hyphomycetes. Bacterial contamination was estimated using Tiara prediction for contigs > 3kb.

| | scaffold | | | | contig | | | | | BUSCO | | | |
|---|---|---|---|---|---|---|---|---|---|---|---|---|---|
| Sample ID | Total sequences | tot. scaffolds | N/L50 | max. length | Total sequences | gaps (%) | tot. Contigs | N/L50 | max. length | GC (%) | complete single reference Genes /BUSCOs | complete-ness (%) n=255 | % bacterial conta-mination |
| 27 | 22060793 | 19650 | 1800/2517 | 52504 | 22057543 | 0.015 | 19687 | 1821/2503 | 52504 | 0.4355 ± 0.0919 | 33 | 13 | 83.4 |
| 28 | 33600897 | 34291 | 3910/1573 | 229644 | 33596797 | 0.012 | 34332 | 3939/1570 | 229644 | 0.4234 ± 0.0882 | 28 | 10.9 | 81.1 |
| 29 | 57908672 | 55945 | 6103/1715 | 148011 | 57900772 | 0.014 | 56024 | 6152/1708 | 148011 | 0.4199 ± 0.0836 | 25 | 9.8 | 89.3 |
| 30 | 28048574 | 31771 | 4170/1269 | 61679 | 28045554 | 0.011 | 31803 | 4194/1265 | 61679 | 0.4154 ± 0.0863 | 23 | 9.1 | 79.4 |
| 31 | 11750779 | 13616 | 1679/1116 | 46857 | 11749209 | 0.013 | 13638 | 1697/1113 | 46857 | 0.4385 ± 0.0879 | 14 | 5.5 | 89.9 |
| 32 | 45693586 | 37148 | 545/16442 | 167801 | 45682586 | 0.024 | 37258 | 587/15462 | 138988 | 0.4444 ± 0.0891 | 205 | 80.4 | 17.0 |
| 33 | 11616418 | 5298 | 119/14562 | 252092 | 11612949 | 0.03 | 5346 | 139/13555 | 250564 | 0.5164 ± 0.0800 | 28 | 11.8 | 51.5 |
| 34 | 26317197 | 3668 | 111/75088 | 272418 | 26313497 | 0.014 | 3705 | 116/69334 | 272418 | 0.4901 ± 0.0688 | 251 | 100 | 0.4 |
| 35 | 29497585 | 33534 | 4694/1285 | 49446 | 29491285 | 0.021 | 33597 | 4730/1281 | 49446 | 0.4270 ± 0.0836 | 34 | 15.3 | 62.8 |
| 36 | 18154988 | 23123 | 3868/955 | 39293 | 18152878 | 0.012 | 23145 | 3891/953 | 39293 | 0.4183 ± 0.0778 | 12 | 5.5 | 86.3 |
| 37 | 47305764 | 45708 | 4002/2054 | 150430 | 47290064 | 0.033 | 45865 | 4116/2028 | 150430 | 0.4237 ± 0.0866 | 25 | 14.1 | 80.2 |

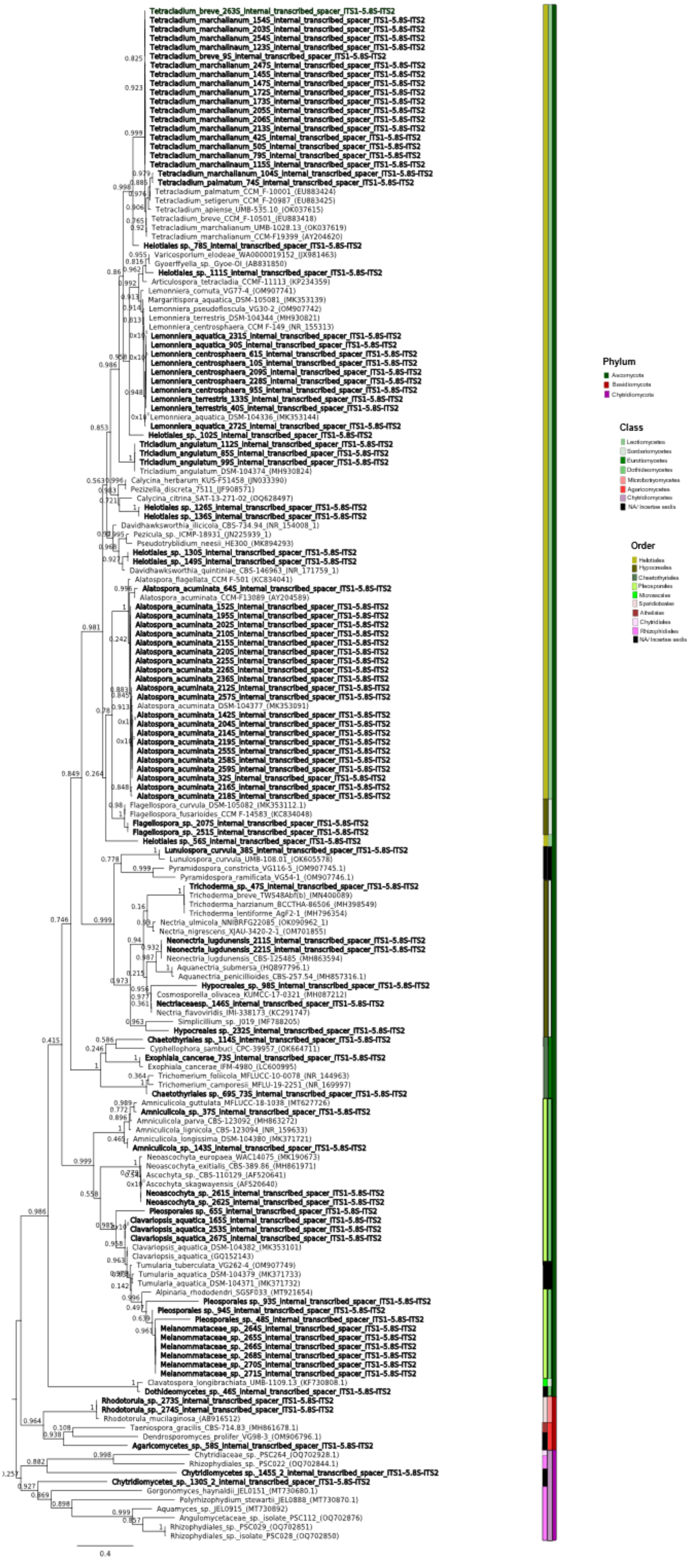

Tetracladium_breve_263S_internal_transcribed_spacer_ITS1-5.8S-ITS2
Tetracladium_marchalianum_154S_internal_transcribed_spacer_ITS1-5.8S-ITS2
Tetracladium_marchalianum_203S_internal_transcribed_spacer_ITS1-5.8S-ITS2
Tetracladium_marchalianum_254S_internal_transcribed_spacer_ITS1-5.8S-ITS2
Tetracladium_marchalinaum_123S_internal_transcribed_spacer_ITS1-5.8S-ITS2
Tetracladium_breve_9S_internal_transcribed_spacer_ITS1-5.8S-ITS2
Tetracladium_marchalianum_247S_internal_transcribed_spacer_ITS1-5.8S-ITS2
Tetracladium_marchalianum_145S_internal_transcribed_spacer_ITS1-5.8S-ITS2
Tetracladium_marchalianum_147S_internal_transcribed_spacer_ITS1-5.8S-ITS2
Tetracladium_marchalianum_172S_internal_transcribed_spacer_ITS1-5.8S-ITS2
Tetracladium_marchalianum_173S_internal_transcribed_spacer_ITS1-5.8S-ITS2
Tetracladium_marchalianum_205S_internal_transcribed_spacer_ITS1-5.8S-ITS2
Tetracladium_marchalianum_206S_internal_transcribed_spacer_ITS1-5.8S-ITS2
Tetracladium_marchalianum_213S_internal_transcribed_spacer_ITS1-5.8S-ITS2
Tetracladium_marchalianum_42S_internal_transcribed_spacer_ITS1-5.8S-ITS2
Tetracladium_marchalianum_50S_internal_transcribed_spacer_ITS1-5.8S-ITS2
Tetracladium_marchalianum_79S_internal_transcribed_spacer_ITS1-5.8S-ITS2
Tetracladium_marchalinaum_115S_internal_transcribed_spacer_ITS1-5.8S-ITS2
Tetracladium_marchalianum_104S_internal_transcribed_spacer_ITS1-5.8S-ITS2
Tetracladium_palmatum_74S_internal_transcribed_spacer_ITS1-5.8S-ITS2
Tetracladium_palmatum_CCM_F-10001_(EU883424)
Tetracladium_setigerum_CCM_F-20987_(EU883425)
Tetracladium_apiense_UMB-535.10_(OK037615)
Tetracladium_breve_CCM_F-10501_(EU883418)
Tetracladium_marchalianum_UMB-1028.13_(OK037619)
Tetracladium_marchalianum_CCM-F19399_(AY204620)
Helotiales sp._78S_internal_transcribed_spacer_ITS1-5.8S-ITS2
Varicosporium_elodeae_WA0000019152_(JX981463)
Gyoerffyella_sp._Gyoe-OI_(AB831850)
Helotiales_sp._111S_internal_transcribed_spacer_ITS1-5.8S-ITS2
Articulospora_tetracladia_CCMF-11113_(KP234359)
Lemonniera_cornuta_VG77-4_(OM907741)
Margaritispora_aquatica_DSM-105081_(MK353139)
Lemonniera_pseudofloscula_VG30-2_(OM907742)
Lemonniera_terrestris_DSM-104344_(MH930821)
Lemonniera_centrosphaera_CCM F-149_(NR_155313)
Lemonniera_aquatica_231S_internal_transcribed_spacer_ITS1-5.8S-ITS2
Lemonniera_aquatica_90S_internal_transcribed_spacer_ITS1-5.8S-ITS2
Lemonniera_centrosphaera_61S_internal_transcribed_spacer_ITS1-5.8S-ITS2
Lemonniera_centrosphaera_10S_internal_transcribed_spacer_ITS1-5.8S-ITS2
Lemonniera_centrosphaera_209S_internal_transcribed_spacer_ITS1-5.8S-ITS2
Lemonniera_centrosphaera_228S_internal_transcribed_spacer_ITS1-5.8S-ITS2
Lemonniera_centrosphaera_95S_internal_transcribed_spacer_ITS1-5.8S-ITS2
Lemonniera_terrestris_133S_internal_transcribed_spacer_ITS1-5.8S-ITS2
Lemonniera_terrestris_40S_internal_transcribed_spacer_ITS1-5.8S-ITS2
Lemonniera_aquatica_DSM-104336_(MK353144)
Lemonniera_aquatica_272S_internal_transcribed_spacer_ITS1-5.8S-ITS2
Helotiales_sp._102S_internal_transcribed_spacer_ITS1-5.8S-ITS2
Tricladium_angulatum_112S_internal_transcribed_spacer_ITS1-5.8S-ITS2
Tricladium_angulatum_85S_internal_transcribed_spacer_ITS1-5.8S-ITS2
Tricladium_angulatum_99S_internal_transcribed_spacer_ITS1-5.8S-ITS2
Tricladium_angulatum_DSM-104374_(MH930824)
Calycina_herbarum_KUS-F51458_(JN033390)
Pezizella_discreta_7511_(JF908571)
Calycina_citrina_SAT-13-271-02_(OQ628497)
Helotiales sp._126S_internal_transcribed_spacer_ITS1-5.8S-ITS2
Helotiales sp._136S_internal_transcribed_spacer_ITS1-5.8S-ITS2
Davidhawksworthia_ilicicola_CBS-734.94_(NR_154008_1)
Pezicula_sp._ICMP-18931_(JN225939_1)
Pseudotryblidium_neesii_HE300_(MK894293)
Helotiales_sp._130S_internal_transcribed_spacer_ITS1-5.8S-ITS2
Helotiales_sp._149S_internal_transcribed_spacer_ITS1-5.8S-ITS2
Davidhawksworthia_quintiniae_CBS-146963_(NR_171759_1)
Alatospora_flagellata_CCM F-501 (KC834041)
Alatospora_acuminata_64S_internal_transcribed_spacer_ITS1-5.8S-ITS2
Alatospora_acuminata_CCM-F13089_(AY204589)
Alatospora_acuminata_152S_internal_transcribed_spacer_ITS1-5.8S-ITS2
Alatospora_acuminata_195S_internal_transcribed_spacer_ITS1-5.8S-ITS2
Alatospora_acuminata_202S_internal_transcribed_spacer_ITS1-5.8S-ITS2
Alatospora_acuminata_210S_internal_transcribed_spacer_ITS1-5.8S-ITS2
Alatospora_acuminata_215S_internal_transcribed_spacer_ITS1-5.8S-ITS2
Alatospora_acuminata_220S_internal_transcribed_spacer_ITS1-5.8S-ITS2
Alatospora_acuminata_225S_internal_transcribed_spacer_ITS1-5.8S-ITS2
Alatospora_acuminata_226S_internal_transcribed_spacer_ITS1-5.8S-ITS2
Alatospora_acuminata_236S_internal_transcribed_spacer_ITS1-5.8S-ITS2
Alatospora_acuminata_212S_internal_transcribed_spacer_ITS1-5.8S-ITS2
Alatospora_acuminata_257S_internal_transcribed_spacer_ITS1-5.8S-ITS2
Alatospora_acuminata_DSM-104377_(MK353091)
Alatospora_acuminata_142S_internal_transcribed_spacer_ITS1-5.8S-ITS2
Alatospora_acuminata_204S_internal_transcribed_spacer_ITS1-5.8S-ITS2
Alatospora_acuminata_214S_internal_transcribed_spacer_ITS1-5.8S-ITS2
Alatospora_acuminata_219S_internal_transcribed_spacer_ITS1-5.8S-ITS2
Alatospora_acuminata_255S_internal_transcribed_spacer_ITS1-5.8S-ITS2
Alatospora_acuminata_258S_internal_transcribed_spacer_ITS1-5.8S-ITS2
Alatospora_acuminata_259S_internal_transcribed_spacer_ITS1-5.8S-ITS2
Alatospora_acuminata_32S_internal_transcribed_spacer_ITS1-5.8S-ITS2
Alatospora_acuminata_216S_internal_transcribed_spacer_ITS1-5.8S-ITS2
Alatospora_acuminata_218S_internal_transcribed_spacer_ITS1-5.8S-ITS2
Flagellospora_curvula_DSM-105082_(MK353112.1)
Flagellospora_fusarioides_CCM F-14583_(KC834048)
Flagellospora_sp._207S_internal_transcribed_spacer_ITS1-5.8S-ITS2
Flagellospora_sp._251S_internal_transcribed_spacer_ITS1-5.8S-ITS2
Helotiales sp._56S_internal_transcribed_spacer_ITS1-5.8S-ITS2
Lunulospora_curvula_38S_internal_transcribed_spacer_ITS1-5.8S-ITS2
Lunulospora_curvula_UMB-108.01_(OK605578)
Pyramidospora_constricta_VG116-5_(OM907745.1)
Pyramidospora_ramificata_VG54-1_(OM907746.1)
Trichoderma_sp._47S_internal_transcribed_spacer_ITS1-5.8S-ITS2
Trichoderma_breve_TW548Abf(b)_(MN400089)
Trichoderma_harzianum_BCCTHA-86506_(MH398549)
Trichoderma_lentiforme_AgF2-1_(MH796354)
Nectria_ulmicola_NNIBRFG22085_(OK090962_1)
Nectria_nigrescens_XJAU-3420-2-1_(OM701855)
Neonectria_lugdunensis_211S_internal_transcribed_spacer_ITS1-5.8S-ITS2
Neonectria_lugdunensis_221S_internal_transcribed_spacer_ITS1-5.8S-ITS2
Neonectria_lugdunensis_CBS-125485_(MH863594)
Aquanectria_submersa_(HQ897796.1)
Aquanectria_penicillioides_CBS-257.54_(MH857316.1)
Hypocreales_sp._98S_internal_transcribed_spacer_ITS1-5.8S-ITS2
Cosmosporella_olivacea_KUMCC-17-0321_(MH087212)
Nectriaceaesp._146S_internal_transcribed_spacer_ITS1-5.8S-ITS2
Nectria_flavoviridis_IMI-338173_(KC291747)
Simplicillium_sp._J019_(MF788205)
Hypocreales sp._232S_internal_transcribed_spacer_ITS1-5.8S-ITS2
Chaetothyriales sp._114S_internal_transcribed_spacer_ITS1-5.8S-ITS2
Cyphellophora_sambuci_CPC-39957_(OK664711)
Exophiala_cancerae_73S_internal_transcribed_spacer_ITS1-5.8S-ITS2
Exophiala_cancerae_IFM-4980_(LC600995)
Trichomerium_foliicola_MFLUCC-10-0078_(NR_144963)
Trichomerium_camporesii_MFLU-19-2251_(NR_169997)
Chaetothyriales sp._69S_73S_internal_transcribed_spacer_ITS1-5.8S-ITS2
Amniculicola_guttulata_MFLUCC-18-1038_(MT627726)
Amniculicola_sp._37S_internal_transcribed_spacer_ITS1-5.8S-ITS2
Amniculicola_parva_CBS-123092_(MH863272)
Amniculicola_lignicola_CBS-123094_(NR_159633)
Amniculicola_longissima_DSM-104380_(MK371721)
Amniculicola_sp._143S_internal_transcribed_spacer_ITS1-5.8S-ITS2
Neoascochyta_europaea_WAC14075_(MK190673)
Neoascochyta_exitialis_CBS-389.86_(MH861971)
Ascochyta_sp._CBS-110129_(AF520641)
Ascochyta_skagwayensis_(AF520640)
Neoascochyta_sp._261S_internal_transcribed_spacer_ITS1-5.8S-ITS2
Neoascochyta_sp._262S_internal_transcribed_spacer_ITS1-5.8S-ITS2
Pleosporales sp._65S_internal_transcribed_spacer_ITS1-5.8S-ITS2
Clavariopsis_aquatica_165S_internal_transcribed_spacer_ITS1-5.8S-ITS2
Clavariopsis_aquatica_253S_internal_transcribed_spacer_ITS1-5.8S-ITS2
Clavariopsis_aquatica_267S_internal_transcribed_spacer_ITS1-5.8S-ITS2
Clavariopsis_aquatica_DSM-104382_(MK353101)
Clavariopsis_aquatica_(GQ152143)
Tumularia_tuberculata_VG262-4_(OM907749)
Tumularia_aquatica_DSM-104379_(MK371733)
Tumularia_aquatica_DSM-104371_(MK371732)
Alpinaria_rhododendri_SGSF033_(MT921654)
Pleosporales sp._93S_internal_transcribed_spacer_ITS1-5.8S-ITS2
Pleosporales sp._94S_internal_transcribed_spacer_ITS1-5.8S-ITS2
Pleosporales_sp._48S_internal_transcribed_spacer_ITS1-5.8S-ITS2
Melanommataceae_sp._264S_internal_transcribed_spacer_ITS1-5.8S-ITS2
Melanommataceae_sp._265S_internal_transcribed_spacer_ITS1-5.8S-ITS2
Melanommataceae_sp._266S_internal_transcribed_spacer_ITS1-5.8S-ITS2
Melanommataceae_sp._268S_internal_transcribed_spacer_ITS1-5.8S-ITS2
Melanommataceae_sp._270S_internal_transcribed_spacer_ITS1-5.8S-ITS2
Melanommataceae_sp._271S_internal_transcribed_spacer_ITS1-5.8S-ITS2
Clavatospora_longibrachiata_UMB-1109.13_(KF730808.1)
Dothideomycetes_sp._46S_internal_transcribed_spacer_ITS1-5.8S-ITS2
Rhodotorula_sp._273S_internal_transcribed_spacer_ITS1-5.8S-ITS2
Rhodotorula_sp._274S_internal_transcribed_spacer_ITS1-5.8S-ITS2
Rhodotorula_mucilaginosa_(AB916512)
Taeniospora_gracilis_CBS-714.83_(MH861678.1)
Dendrosporomyces_prolifer_VG98-3_(OM906796.1)
Agaricomycetes_sp._58S_internal_transcribed_spacer_ITS1-5.8S-ITS2
Chytridiaceae_sp._PSC264_(OQ702928.1)
Rhizophydiales_sp._PSC022_(OQ702844.1)
Chytridiomycetes sp._145S_2_internal_transcribed_spacer_ITS1-5.8S-ITS2
Chytridiomycetes_sp._130S_2_internal_transcribed_spacer_ITS1-5.8S-ITS2
Gorgonomyces_haynaldii_JEL0151_(MT730680.1)
Polyrhizophydium_stewartii_JEL0888_(MT730870.1)
Aquamyces_sp._JEL0915_(MT730892)
Angulomycetaceae_sp._isolate_PSC112_(OQ702876)
Rhizophydiales_sp._PSC029_(OQ702851)
Rhizophydiales_sp._isolate_PSC028_(OQ702850)
0.4
Phylum
Ascomycota
Basidiomycota
Chytridiomycota
Class
Order

**Figure S1:** Maximum likelihood phylogenetic tree based on ITS1-5.8S-ITS2 region. The tree was rooted with the Chytridiomycota branch (sequences PP384258 and PP386607). Chytridiomycota sequences were obtained from amplicon sequencing results of samples 130S and 145S. Taxa in bold represent species obtained from environmental samples with the described single cell dissection workflow.

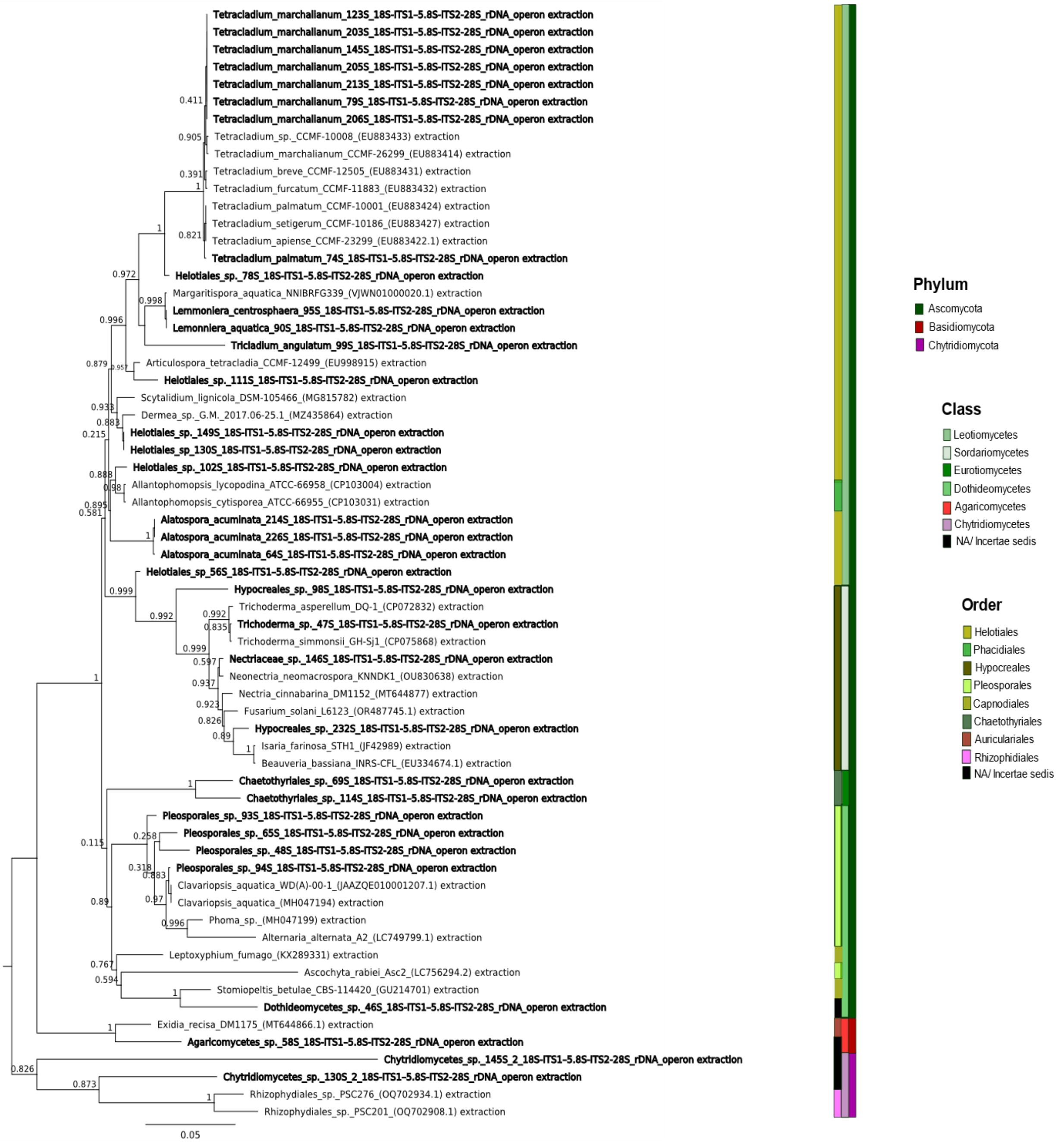


**Figure S2:** Maximum likelihood phylogenetic tree based on the extracted SSU region from the FRO (SSU- ITS1-5.8S-ITS2-LSU) region alignment. The tree was rooted with the Chytridiomycota branch (SSU region of extracted sequences PP384258 and PP386607). Chytridiomycota sequences were obtained from amplicon sequencing results of samples 130S and 145S. Taxa in bold represent species obtained from environmental samples with the described single cell dissection workflow.

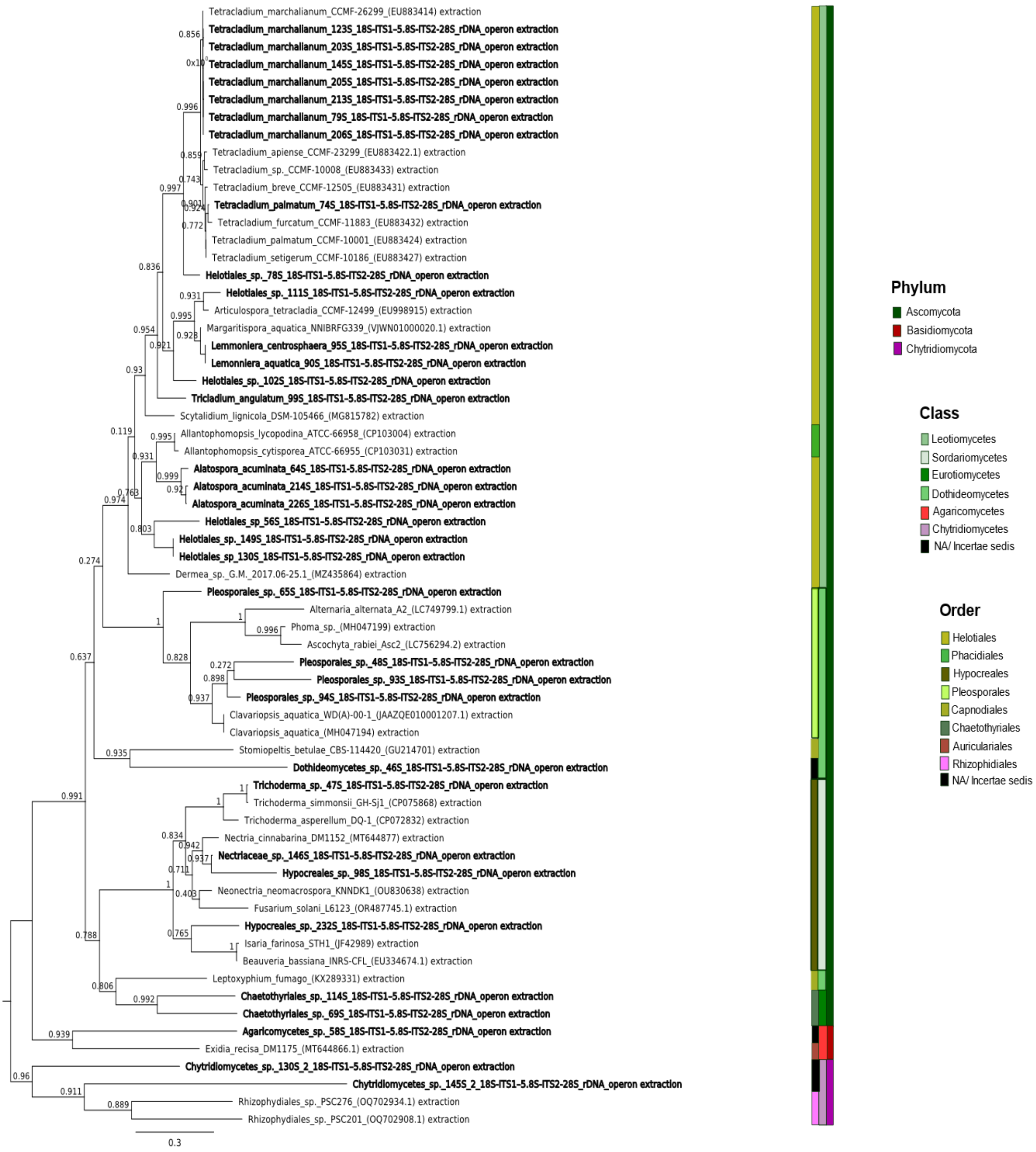


**Figure S3:** Maximum likelihood phylogenetic tree based on the extracted ITS1-5.8S-ITS2 region from the FRO (SSU- ITS1-5.8S-ITS2-LSU) region alignment. The tree was rooted with the Chytridiomycota branch (ITS region of extracted sequences PP384258 and PP386607). Chytridiomycota sequences were obtained from amplicon sequencing results of samples 130S and 145S. Taxa in bold represent species obtained from environmental samples with the described single cell dissection workflow.

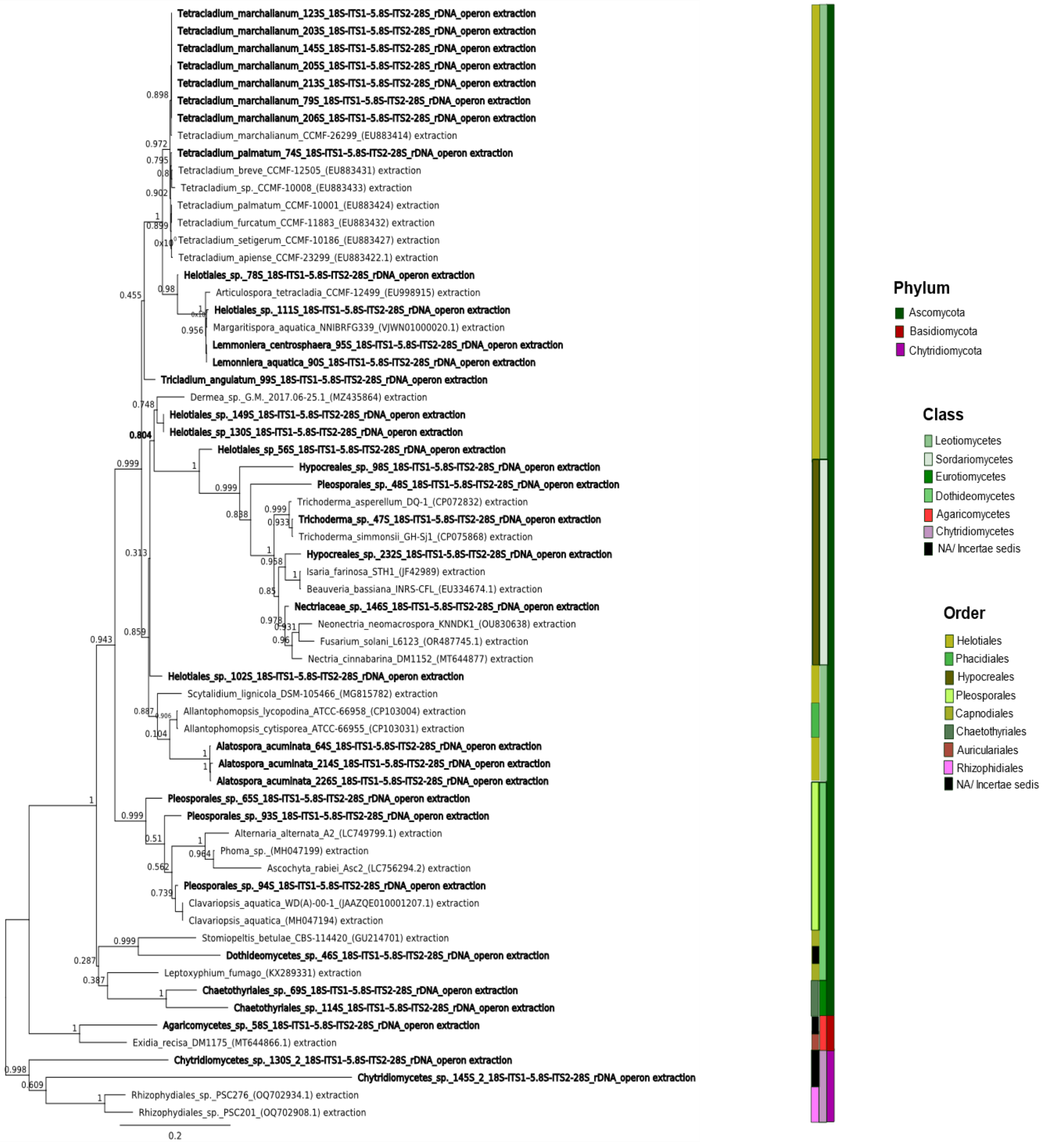


**Figure S4:** Maximum likelihood phylogenetic tree based on the extracted LSU region from the FRO (SSU- ITS1-5.8S-ITS2-LSU) region alignment. The tree was rooted with the Chytridiomycota branch (LSU region of extracted sequences PP384258 and PP386607). Chytridiomycota sequences were obtained from amplicon sequencing results of samples 130S and 145S. Taxa in bold represent species obtained from environmental samples with the described single cell dissection workflow.
