## Supplementary material for "Exploring environmental microfungal diversity through serial single cell screening": Detailed Protocol

### Detailed protocol for Fungal single-cell Laser microdissection (LMD), Whole Genomic Amplification (WGA), and long amplicon sequencing

1. **Sample collection:** Collect plant litter in varying stages of decomposition to sterile plastic bags **(A)**. Alternatively, collect habitat water **(B)**.

2. **Preparation of sporulation assays (A)**: Rinse the leaves with deionized water to remove small debris and macroinvertebrates. Then, insert the leaves in a 200 ml Erlenmeyer flask, submerge them in deionized water, and constantly aerate for 48-72h using an oxygen pump. Plug air inlets with cotton to prevent contamination.

3. **Preparation for laser microdissection:** UV-irradiate a slide holder, collector tube holder, and collection tubes using a UV-crosslinker for 30 minutes at 11.7 J/cm2. UV-irradiate upward-facing PPS-Membrane (4.0 µm) frame slides using a UV-Crosslinker for 30 minutes at 11.7 J/cm2.

   For **(A)**, if the number of cells required is low **(A1)**, transfer 200 µl of the water sample from the Erlenmeyer flasks to the UV-irradiated PPS membrane. Slightly dry samples using a laboratory oven (27ºC, 15min). Stain the membrane with 3-5 drops of calcofluor white 1% (w/v) and incubate it in the dark for 15 min. If the number of cells required is high **(A2)**, filter 20ml of water sample from the Erlenmeyer flasks through a 5.0 µm polycarbonate membrane filter. Stain the filter with 3-5 drops of calcofluor white 1% (w/v), incubate in the dark for 15 min, and place in the UV-irradiated PPS membrane.

   For **(B)**, sift the water sample through a 100 µm followed by a 25 µm sieve and refilter 100 ml of the filtrate through a 5 µm filter. Repeat the process with the remaining initial filtrate volume. Rewet both filters in 5 mL deionized water and re-filter through a 5.0 µm polycarbonate membrane filter. Stain the filter with 3-5 drops of calcofluor white 1% (w/v), incubate it in the dark for 15 min, and place it on the UV-irradiated PPS membrane.

4. **Laser microdissection:** Calibrate the microscope and perform an overview scan to find the fungal cells of interest using the microscope's bright field or fluorescence feature. Mark fungal cells for laser microdissection. For every cutting session, place four 0.2 ml tubes or eight flat collection caps in the collector tube holder.

   To perform cell microdissection, the following parameters are recommended:

| Sample Type | Power | Aperture | Speed | Specimen balance | Offset | Head current (%) | Pulse Frequency (Hz) |
|---|---|---|---|---|---|---|---|
| A1 | 28-37 | 14-20 | 4-6 | 18-24 | 50 | 100 | 120 |
| A2 | 20-27 | 2-9 | 5-8 | 13-20 | 50 | 100 | 120 |
| B | 20-27 | 2-9 | 5-8 | 13-20 | 50 | 100 | 120 |

*Note:* At lower magnification (10x), the power settings can be further reduced.

   After dissection, inspect the collection caps to count the successfully dissected target cells. Transfer the collection tube holder to a disinfected and UV-decontaminated PCR workstation for further processing.

5. **Cell preparation**: Add 6 µl of Advanced storage buffer (REPLI-g Advanced DNA Single Cell Kit – Qiagen) to each cell sample. Place the 8-cap strips into the tubes/plates, and centrifuge in a microcentrifuge (3000rpm; 3min).

   Optional stopping point: At this point you can store samples at -20ºC.

6. **Enzymatic lysis**: Add 5 units of Zymolyase (Zymo Research, CA, USA) to each reaction tube. Briefly centrifuge the samples using a microcentrifuge and incubate for 60 min at 37 °C. Deactivate Zymolyase by incubating samples at 65 °C for 10 min.

7. **Alkaline lysis:** Prepare the D2 solution from the REPLI-g Advanced DNA Single Cell Kit according to the manufacturer's instructions. Add 3 µl of D2 to each reaction tube and briefly centrifuge. Incubate samples at 65 °C for 10 min. Add 3 µl of Stop solution to each reaction tube to stop the reaction. Mix and centrifuge (3000rpm; 1min).

   Optional stopping point: At this point you can store samples overnight at 4ºC .

8. **Multiple displacement amplification (MDA)**: Perform the MDA reaction according to the instructions in REPLI-g Advanced DNA Single Cell Kit (Qiagen), with an incubation period of 4 h at 30 °C. Check the successful amplification by gel electrophoresis (3 µl MDA product in 0.8% w/v agarose; 90min; 80V).

9. **Dilution of MDA products**: Dilute 1 µl of MDA product in 99 µl of Nuclease-free water (1:100 dilution).

10. **Optional: Amplification of the Fungal ITS1-5.8S-ITS2 region:** Prepare a PCR reaction mix (GoTaq® Green Master Mix, Promega) without template as indicated and dispatch 22 µl of reaction mix per reaction tube. Add 3 µl of MDA diluted product per reaction, mix carefully, and spin the reaction down. Cycle the reactions according to the conditions below.

**Primers used in this amplification:**

| Primer name | Sequence |
|---|---|
| **ITS4-Ad** | GTC TCG TGG GCT CGG AAT CCT CCG CTT ATT GAT ATG C |
| **ITS5-Ad** | TCG GCA GCG TCT TGG AAG TAA AAG TCG TAA CAA GG |

**Amplicon PCR Reaction mix (ITS5/ITS4):**

| Reagent | Stock conc. | Final conc. | Vol. For 1 Reaction (µl) |
|---|---|---|---|
| | | | For 25 µl |
| **Nuclease free $H_2O$** | | | 7 |
| **Primer ITS5-Ad fwd** | 10 µM | 0.5 µM | 1.25 |
| **Primer ITS4-Ad rev** | 10 µM | 0.5 µM | 1.25 |
| **GoTaq® Green Master Mix** | 2X | 1X | 12.5 |
| **MDA product (1:100 Diluted)** | 50 - 200 ng | 0.5 - 2 ng | 3 |
| **Total volume** | - | - | 25 |

**PCR cycling conditions (ITS5/ITS4):**

| Step | Temp. | Time | Cycles |
|---|---|---|---|
| **Activation** | 94ºC | 2 min | |
| **Denaturation** | 94ºC | 1 min | |
| **Annealing** | 55ºC | 1 min | x 35 |
| **Elongation** | 72ºC | 2 min | |
| **Final elongation** | 72ºC | 10 min | |
| **Hold** | 4°C | ∞ | |

11. **Optional: Confirmation of ITS1-5.8S-ITS2 region (0.5-1.5 kb) PCR success:** Perform gel electrophoresis, adding 3 µl of PCR product to a 0.8% agarose gel.

12. **Amplification of the Fungal full rDNA operon (NS1short/RCA95m):** Mix all reagents except the DNA template and dispatch 28 µl per reaction well. Add 2 µl of diluted MDA product and cycle the reactions as indicated below.

**Primers used in this amplification:**

| Primer name | Sequence |
|---|---|
| **NS1-short-Ad** | TCG TCG GCA GCG TCT TCA GTA GTC ATA TGC TTG TC |
| **RCA95m-Ad** | GTC TCG TGG GCT CGG AAC TAT GTT TTA ATT AGA CAG TCA G |

**Amplicon PCR Reaction mix (NS1short/RCA95m):**

| Reagent | Stock conc. | Final conc. | Vol. For 1 Reaction (µl) |
|---|---|---|---|
| | | | For 30 µl |
| **H2O** | | | 11.5 |
| **Primer NS1 Short fwd** | 10 µM | 0.25 µM | 0.75 |
| **Primer RCA95m rev** | 10 µM | 0.25 µM | 0.75 |
| **PrimeSTAR GXL Master Mix (Takara Bio)** | 2X | 1X | 15 |
| **MDA product (1:100 Diluted)** | 50 - 200 ng | 0.5 - 2 ng | 2 |
| **Total volume** | | | 30 |

**PCR cycling conditions (NS1short/RCA95m):**

| Step | Temp. | Time | Cycles |
|---|---|---|---|
| **Activation** | 98ºC | 1 min | |
| **Denaturation** | 98ºC | 10 sec | x 35 |
| **Annealing** | 55ºC | 15 sec | |
| **Elongation** | 68ºC | 2:30 min | |
| **Hold** | 12°C | ∞ | |

13. **Confirmation of the full rDNA operon (NS1short/RCA95m; 4-6 kbp) PCR success:** Perform gel electrophoresis, adding 3 µl of PCR product to a 0.8% agarose gel.
14. **PCR products clean up:** Purify successful PCR products using 0.8 v/v of AMPure XP beads (Beckman Coulter) according to the manufacturer's instructions.
15. **Confirmation of the full rDNA operon PCR clean-up success:** Perform a gel electrophoresis, adding 3 µl of PCR product to a 0.8% agarose gel. Add a 3 µl of MassRuler High Range DNA Ladder (ThermoScientific™) to the gel to estimate the PCR product concentration. Assign the PCR products to 4 scales (2, 4, 6 and 8) based on the estimated concentration. This is relevant to determine the amount of purified PCR product added to the indexing PCR reaction in the next step.

16. **Indexing PCR:** Prepare the sample indexing scheme (see indexing primers below).

**Index Fwd primers**

| | |
|---|---|
| **for-bc-a** | AAGAAAGTTGTCGGTGTCTTTGTGTCGTCGGCAGCGTC |
| **for-bc-b** | TCGATTCCGTTTGTAGTCGTCTGTTCGTCGGCAGCGTC |
| **for-bc-c** | GAGTCTTGTGTCCCAGTTACCAGGTCGTCGGCAGCGTC |
| **for-bc-d** | TTCGGATTCTATCGTGTTTCCCTATCGTCGGCAGCGTC |
| **for-bc-e** | CTTGTCCAGGGTTTGTGTAACCTTTCGTCGGCAGCGTC |
| **for-bc-f** | TTCTCGCAAAGGCAGAAAGTAGTCTCGTCGGCAGCGTC |
| **for-bc-g** | GTGTTACCGTGGGAATGAATCCTTTCGTCGGCAGCGTC |
| **for-bc-h** | TTCAGGGAACAAACCAAGTTACGTTCGTCGGCAGCGTC |
| **for-bc-i** | AACTAGGCACAGCGAGTCTTGGTTTCGTCGGCAGCGTC |
| **for-bc-j** | AAGCGTTGAAACCTTTGTCCTCTCTCGTCGGCAGCGTC |
| **for-bc-k** | GTTTCATCTATCGGAGGGAATGGATCGTCGGCAGCGTC |
| **for-bc-l** | CAGGTAGAAAGAAGCAGAATCGGATCGTCGGCAGCGTC |

| Index Rev primers | |
|---|---|
| **rev-bc-1** | TGTGTTGAGACCACACAGGCCTCAGTCTCGTGGGCTCGG |
| **rev-bc-2** | GTCTGTCGCCATGGAAAGTCAACTGTCTCGTGGGCTCGG |
| **rev-bc-3** | TTGCTACGGTTGACCATGCAGTTAGTCTCGTGGGCTCGG |
| **rev-bc-4** | AACTTGAGGTATCGTATATTCAATGTCTCGTGGGCTCGG |
| **rev-bc-5** | GCAGGTGGGCATCCGGACCGATATGTCTCGTGGGCTCGG |
| **rev-bc-6** | CAGAGCTGACCCTCCAGATATTTGGTCTCGTGGGCTCGG |
| **rev-bc-7** | TCTTAGTGTATGAGCTCGCTCACCGTCTCGTGGGCTCGG |
| **rev-bc-8** | CCCTGGGACGTAGGAATCCACGCCGTCTCGTGGGCTCGG |
| **rev-bc-9** | TGTTGCGAACGGGACCTGCCTAGCGTCTCGTGGGCTCGG |
| **rev-bc-10** | ACACCTTTACATAGGCCGCCATCTGTCTCGTGGGCTCGG |
| **rev-bc-11** | GACCTTAGTCACATGGTAGTCTAAGTCTCGTGGGCTCGG |
| **rev-bc-12** | GTTCGGATGCAATATGGTTCACTGGTCTCGTGGGCTCGG |
| **rev-bc-13** | TAGCAGAAGTCCCTGTAAGACCATGTCTCGTGGGCTCGG |
| **rev-bc-14** | GATTCTGATTACTCTATTCGCCAGGTCTCGTGGGCTCGG |
| **rev-bc-15** | GGAATAATACCATTGAAGTAGCACGTCTCGTGGGCTCGG |
| **rev-bc-16** | GGGTCCCTCTACTCATTTAGCATGGTCTCGTGGGCTCGG |

Combine each sample with a unique index combination, e.g., use a plate layout as indicated below, with forward index primers on the plate y-axis and reverse index primers on the x-axis.

| | rev-bc-1 | rev-bc-2 | rev-bc-3 | rev-bc-4 | rev-bc-5 | 6 | 7 | 8 | 9 | 10 | 11 | 12 |
|---|---|---|---|---|---|---|---|---|---|---|---|---|
| for-bc-a | S1 | S4 | S7 | S10 | S13 | | | | | | | |
| for-bc-b | S2 | S5 | S8 | S11 | S14 | | | | | | | |
| for-bc-c | S3 | S6 | S9 | S12 | S15 | | | | | | | |
| D | | | | | | | | | | | | |
| E | | | | | | | | | | | | |
| F | | | | | | | | | | | | |
| G | | | | | | | | | | | | |
| H | | | | | | | | | | | | |
| PLATE 1 - Index Primer fwd – (right- left – 8 channel) | | | | | | | | | | | | |

For the indexing PCR, prepare 4 Mixes (see step 15, according to the scale of the PCR product concentration 2,4,6,8) with the corresponding amount of PrimeSTAR GXL (Takara Bio) master mix and water, dispatching 29.6-35.6 µl per reaction well, depending on the chosen Mix. Add indexing primers and template separately. Mix the reactions by pipetting and spin them down. Cycle the reactions according to the protocol below.

**Indexing PCR Reaction mix:**

| Reagent | Stock conc. | Final conc. | Vol. 1 Reaction (µl) | | | |
|---|---|---|---|---|---|---|
| | | | For 40 µl | | | |
| PrimeSTAR GXL Master Mix | 2X | 1X | 20 | | | |
| $H_2O$ | | | 9.6 | 11.6 | 13.6 | 15.6 |
| Primer index fwd | 10 µM | 0.3 µM | 1.2 | | | |
| Primer index rev | 10 µM | 0.3 µM | 1.2 | | | |
| Purified PCR product | Ca. 30-100 ng/ul | 100-400 ng | 2 | 4 | 6 | 8 |
| Total volume | | | 40 | | | |

**Indexing PCR cycling conditions:**

| Step | Temp. | Time | Cycles |
|---|---|---|---|
| Activation | 98ºC | 1 min | |
| Denaturation | 98ºC | 10 sec | |
| Annealing | 55ºC | 15 sec | x 4 |
| Elongation | 68ºC | 6 min | |
| Hold | 12°C | ∞ | |

17. **Confirmation of indexing PCR success**: Perform a gel electrophoresis with 3 µl of indexing PCR product (0.7v/v; 70min;80V).

18. **PCR products clean up**: Purify successful PCR products using 0.8 v/v of AMPure XP beads according to the manufacturer's instructions.

19. **DNA quantification:** Quantify the purified indexing PCR products using a fluorophore bases dsDNA quantification system, following the manufacturer’s instructions. Use the results to calculate the DNA concentration in each of the samples.

20. **Pooling:** Pool PCR products in approximately equimolar quantities in a 1.5ml Eppendorf tube. Note the differences in fragment lengths for molarity calculation.

21. **Pool clean-up:** Purify the sequencing pool using 0.8 v/v of AMPure XP beads per the manufacturer's instructions and quantify the DNA pool using a fluorometric dsDNA quantification assay.

22. **Oxford Nanopore Library preparation:** Use 100-200 fmol of the purified DNA pool for the sequencing adapter ligation. Follow the Oxford Nanopore library preparation protocol of the **ligation sequencing kit**, e.g., SQK-LSK114, with the associated protocol for amplicon sequencing. Perform all steps according to the manufacturer’s protocol.

23. **Sequencing:** Load the final library into your flow cell (e.g., FLO-MIN114) and perform sequencing using your sequencing device (e.g., MinION). You can stop the sequencing run once your desired sequencing depth (e.g., 1,000 – 2,000 reads per sample) is achieved.

24. **Data processing:** After data acquisition, perform the following steps.

    i. Basecall the fast5 sequencing files as fastq files in the high accuracy base-calling mode using an up-to-date basecaller (currently: Dorado).

    ii. Optional: basecall duplex read pairs using Dorado for Q30 reads (Previously ONT’s duplex tools: https://pypi.org/project/duplex-tools/; v0.2.9)

    iii. Process the basecalled FASTQ files using the following bioinformatic pipeline steps following Wurzbacher et al. 2019 (see below) or an equivalent pipeline for consensus generation.

a) sequence length filtering using Biopython (Cock et al. 2009), to exclude reads outside of the expected length range (Wurzbacher et al. 2019)

b) Demultiplexing with Flexbar v2.5 (Dodt, Roehr et al. 2012), to sort sequenced reads of each individual sample according to its unique index

c) Alignment of demultiplexed FASTA files using MAFFT (v7.397) (Katoh and Standley 2013) with the auto-alignment option

d) Clustering in mothur v1.39 (Schloss, Westcott et al. 2009) using the Opticlust algorithm

e) Generation of final consensus sequences for each operational taxonomic unit with Consension ( v1.0, https://microbiology.se/software/consension/).

25. **Optional quality control step for bacterial contamination (required for whole genome sequencing):** Perform a bacterial 16S rRNA quantitative PCR to test the WGA amplified DNA product for bacterial contamination using the 16S V3-V4 primers published by Muyzer et *al*. 1993 (DOI: https://doi.org/10.1128/aem.59.3.695-700.1993):

| Target | Name | Sequence 5`->3` | TA (°C) | Expected Fragment length (bp) |
|---|---|---|---|---|
| V3-V4 | 337f | GAC TCC TAC GGG AGG CWG CAG | 60°C | ~ 200 |
| | 518r | GTA TTA CCG CGG CTG CTG G | | |

i. Prepare a 96-well plate set-up of the qPCR reactions and prepare the sample setup in the qPCR machine.

| | 1 | 2 | 3 | 4 | 5 | 6 | 7 | 8 | 9 | 10 | 11 | 12 |
|---|---|---|---|---|---|---|---|---|---|---|---|---|
| A | Std1 | Std1 | | NTC | NTC | | | | | | | |
| B | Std2 | Std2 | | | | | | | | | | |
| C | Std3 | Std3 | | C1 | C1 | | | | | | | |
| D | Std4 | Std4 | | C2 | C2 | | | | | | | |
| E | Std5 | Std5 | | C3 | C3 | | | | | | | |
| F | Std6 | Std6 | | Cn | Cn | | | | | | | |
| G | Std7 | Std7 | | | | | | | | | | |
| H | Std8 | Std8 | | | | | | | | | | |

*Note: Std = Standard DNA, NTC = negative template control. C1 – Cn: WGA DNA samples*

ii. Thaw all reagents on ice. If thawed, gently mix and spin them down. Prepare primer and template dilutions to the desired concentration.

iii. Perform six to eight serial dilutions of the standard DNA (purified and quantified PCR product of full-length 16S rRNA V1-V9 region of E. *coli*). Perform e.g. a serial 1:10 dilutions resulting in eight tubes with: $0.5 \cdot 10^9 > 0.5 \cdot 10^8 > 0.5 \cdot 10^7 > 0.5 \cdot 10^6 > 0.5 \cdot 10^5 > 0.5 \cdot 10^4 > 0.5 \cdot 10^3 > 0.5 \cdot 10^2$ copies / µl.

iv. Prepare the master mix by adding all reagents in the order of the table below. Keep the solution light - protected using aluminum foil. Each reaction has a final volume of 20 µl.

**Q-PCR Reaction mix (16S rRNA V3-V4):**

| Reagent | Stock conc. | Final conc. | Vol. For 1 Reaction (µl) |
|---|---|---|---|
| | | | For 20 µl |
| **H2O** | | | 6 |
| **Primer 337f** | 10 µM | 0.5 µM | 1 |
| **Primer 518r** | 10 µM | 0.5 µM | 1 |
| **NEB Luna® Universal qPCR Master Mix** | 2X | 1X | 10 |
| **MDA product (1:100 Diluted) /Std** | 50 - 200 ng | 0.5 - 2 ng | 2 |
| **Total volume** | | | 20 |

v. Prepare the master mix by adding all reagents in the order of the table below. Keep the solution light - protected using aluminum foil. Each reaction has a final volume of 20 µl.

i. Distribute the master mix in the corresponding wells of a UV-radiated 96-well plate. Then, add the template and standard DNA before sealing and spinning the plate.

ii. Cycle the qPCR reaction according to the cycling regime indicated below on a Bio-Rad CFX-connect real-time PCR detection system:

**Q-PCR cycling conditions (16S rRNA V3-V4):**

| Step | Temp. | Time | Cycles |
|---|---|---|---|
| **Activation** | 95ºC | 1 min | |
| **Denaturation** | 95ºC | 15 sec | x 35 |
| **Annealing + Elongation** | 60ºC | 40 sec | |
| **Plate read** | | | |
